# Chromatin activity identifies differential gene regulation across human ancestries

**DOI:** 10.1101/2022.11.25.517959

**Authors:** Kade P. Pettie, Maxwell Mumbach, Amanda J. Lea, Julien Ayroles, Howard Y. Chang, Maya Kasowski, Hunter B. Fraser

**Affiliations:** Department of Biology, Stanford University, Stanford, CA; Department of Genetics, Stanford University, Stanford, CA; Department of Biological Sciences, Vanderbilt University, Nashville, TN; Department of Ecology and Evolutionary Biology, Princeton University, Princeton, NJ; Center for Personal Dynamic Regulomes, Stanford University School of Medicine, Stanford, CA; Howard Hughes Medical Institute, Stanford University, Stanford, CA; Sean N. Parker Center for Allergy and Asthma Research, Stanford University School of Medicine, Stanford, CA; Department of Pathology, Stanford University School of Medicine, Stanford, CA

## Abstract

Current evidence suggests that *cis*-regulatory elements controlling gene expression may be the predominant target of natural selection in humans and other species. Detecting selection acting on these elements is critical to understanding evolution but remains challenging because we do not know which mutations will affect gene regulation. To address this, we devised an approach to search for lineage-specific selection on chromatin activity, transcription factor binding, and chromosomal looping—critical steps in transcriptional regulation. Applying this approach to lymphoblastoid cells from 831 individuals of either European or African descent, we find strong signals of differential chromatin activity linked to gene expression differences between ancestries in numerous contexts, but no evidence of functional differences in chromosomal looping. Moreover, we show that enhancers rather than promoters display the strongest signs of selection associated with sites of differential transcription factor binding. Overall, our study indicates that some *cis*-regulatory adaptation may be more easily detected at the level of chromatin than DNA sequence. This work provides a vast resource of genomic interaction data from diverse human populations and establishes a novel selection test that will benefit future study of regulatory evolution in humans and other species.

## Introduction

Identifying the genetic determinants of complex traits is challenging because their contributions are often diluted across many variants of small effect. Single variants of large effect are simpler to identify and have been well-characterized (Shapiro et al. 2004; Tishkoff et al. 2007; Manceau et al. 2011), but genome-wide association studies (GWAS), which test millions of variants for statistical association with a trait, have demonstrated that these large-effect loci are rare. Moreover, the vast majority of trait-associated variants are located in non-coding regions (Maurano et al. 2012; Schaub et al. 2012; Gallagher and Chen-Plotkin 2018; Cano-Gamez and Trynka 2020). For the < 10% of GWAS hits in protein-coding regions, inferences about their evolutionary history and mechanisms of action are often readily available thanks to studies that have focused on these regions. The remaining > 90% of GWAS hits in non-coding regions are thought to affect traits by altering gene expression levels, but causal mechanisms are obscured by a combination of linkage disequilibrium (LD), a genome-wide phenomenon in which nearby variants tend to be inherited together leading to a correlation of their effects (Sohail et al. 2020), and the paucity of information about non-coding relative to coding regions.

Even before the GWAS era variants with highly divergent allele frequencies between populations, measured by estimates of Wright’s fixation index (*F_ST_*) (Weir and Cockerham 1984), were found to be enriched in disease-associated genes (Foll and Gaggiotti 2008). Since then, genome-wide scans using associations of allele frequencies with environmental variables as evidence of natural selection have shown signals of positive selection to be somewhat enriched in coding regions (Coop et al. 2010; Fumagalli et al. 2010; Hancock et al. 2010, 2011a, 2011b), and even more enriched in cis-regulatory elements (Fraser 2013). Overall, the preponderance of non-coding variants implicated in human GWAS is paralleled by a similar trend among human genetic variants involved in local environmental adaptation (Fraser 2013).

Intersecting non-coding GWAS hits with information from assays measuring regulatory activity, such as quantitative trait loci (QTL) for molecular-level traits (mol-QTL), has been effective at pinpointing causal variants and molecular mechanisms underlying complex trait variation (Grubert et al. 2015; Kaplow et al. 2015; Tehranchi et al. 2016, 2019; Greenwald et al. 2019; GTEx Consortium 2020). QTL studies using gene expression as the trait (eQTL) test all variants within a predefined distance (usually one megabase (Mb)) of a gene for an association with that gene’s expression, so each eQTL is linked to a target gene (GTEx Consortium 2020). Since transcription factor (TF) proteins bind gene regulatory elements such as enhancers in a sequence-dependent manner to regulate transcription, eQTL can act by altering a TF’s binding affinity (*i.e.*, one allele has higher binding affinity than the other, termed a bQTL) (Tehranchi et al. 2016). In most cases, increased TF binding is associated with decompaction of chromatin, the DNA-protein complex that packages meters of linear DNA into a nucleus a few microns wide. This opening of the chromatin allows more TFs to bind to previously inaccessible stretches of DNA and to each other in a positive feedback loop of chromatin accessibility. Thus, chromatin accessibility can be used as a proxy for regulatory activity to identify enhancers and their relative activity levels, as is accomplished with Assay for Transpose-Accessible Chromatin (ATAC-seq) (Buenrostro et al. 2015).

Since enhancers operate in three-dimensional space and can contact target gene promoters (*cis* regulation) several Mb away, ATAC-seq and high-throughput methods based on Chromatin Conformation Capture (HiC) (Lieberman-Aiden et al. 2009; Mifsud et al. 2015; Mumbach et al. 2016, 2019) can be combined to identify enhancer-promoter interactions (Waszak et al. 2015; Grubert et al. 2015; Tehranchi et al. 2016, 2019). The activity-by-contact (ABC) model was recently developed to predict enhancer-target gene pairs in a given cell type under the premise that the extent to which an element regulates a gene’s expression depends on its strength as an enhancer (activity level), scaled by how often it is near that gene’s promoter in 3D space (contact frequency) (Fulco et al. 2019). HiChIP, which combines HiC with chromatin immunoprecipitation (ChIP) on a protein of interest, is well-suited to generate input for this model, particularly when performed on the histone modification H3K27ac, a hallmark of active chromatin. Since the end product is paired-end reads from H3K27ac-associated long-range interactions, H3K27ac HiChIP provides a simultaneous measure of activity level and contact frequency without the high sequencing depth and cell number required to generate the all-by-all interaction maps of HiC (Mumbach et al. 2016, 2017). The ABC model has been shown to attain peak performance with chromatin accessibility and HiChIP data as input and outperforms other enhancer target gene prediction methods (Fulco et al. 2019), making it a powerful metric for hypothesis generation about the mechanisms of non-coding GWAS hits (Nasser et al. 2021).

Additional support for the mechanisms and causality of these hits can come from intersecting molecular-level QTL with putative locally adaptive variants (Fraser 2013). However, since selection acts on fitness, its impact may be more directly observable at the level of chromatin activity than at the level of DNA sequence, where it is relatively more diluted (Fig. 1a, left). For example, chromatin activity is a better predictor of TF binding than DNA sequence since we do not fully understand the *cis*-regulatory “code” that governs TF binding (Peng et al. 2019).

**Figure 1.**
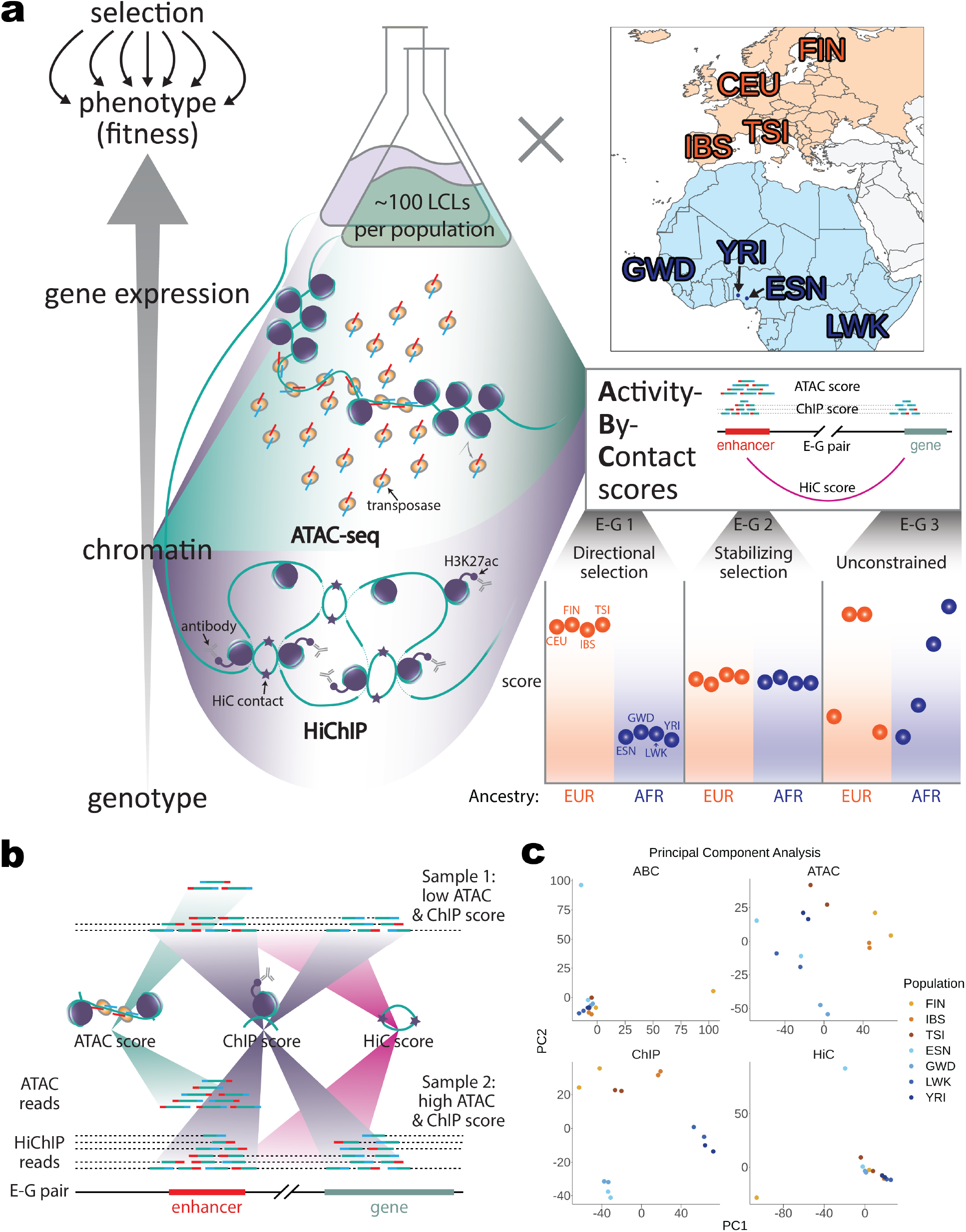
Identification of candidate enhancer-gene pairs under different selection pressures between ancestries. **a)** From left to right: Chromatin is closer than genotype to where selection acts directly. ATAC-seq to measure chromatin accessibility and H3K27ac HiChIP to measure chromatin activity and contact frequency were performed in LCLs from four African populations (blue) and four European populations (orange). Activity-by-contact (ABC) scores were computed to define candidate enhancer-gene pairs and ATAC, ChIP, and HiC scores representing each ABC component were computed for each enhancer-gene pair. Examples of ATAC and HiChIP sequencing read pileups are shown contributing to each component score. Enhancer-gene pairs with low within- and high between-continental ancestry score variance are suggestive of directional selection, shown here for enhancer-gene pair 1. **b)** A more detailed depiction of sequencing reads’ contributions to each ABC component score is shown for a hypothetical enhancer-gene pair where sample 2 has higher ATAC and ChIP scores, but an equivalent HiC score relative to sample 1. ATAC scores represent the ATAC-seq signal at the enhancer (teal gradients). ChIP scores estimate the enhancer-gene pair’s collective H3K27ac signal as the geometric mean of total HiChIP signal at the enhancer and gene promoter (also known as the vanilla coverage square root (VC-sqrt)). Dotted lines extending to the depicted region border connect to reads whose other paired end aligns elsewhere in the genome but contributes to the HiChIP score (purple gradient). HiC scores estimate the contact frequency of the enhancer and promoter independent of H3K27ac levels by dividing the HiChIP signal from read pairs specifically connecting the enhancer and promoter (magenta gradient) by the VC-sqrt. **c)** Principal component analysis results from all enhancer-gene pairs are shown for each score type. Both replicates are shown for each population. CEU is an outlier (Fig. S4-7) and is thus excluded in these PCAs and in downstream analyses. Abbreviations: ATAC-seq = Assay for Transpose Accessible Chromatin with sequencing; E = enhancer; G = gene; H3K27ac = histone 3 lysine 27 acetylation; LCLs = Lymphoblastoid Cell Lines; CEU = Utah residents with North European ancestry; ESN = Esan; FIN = Finnish; GWD = Gambian; IBS = Iberian; LWK = Luhya; TSI = Tuscan; YRI = Yoruban; AFR = African; EUR = European; PC = principal component.

Studies in primates have suggested that directional selection may have contributed to differences in chromatin activity that distinguish each species (Edsall et al. 2019). For example, sites with decreased chromatin accessibility in human relative to chimpanzee and rhesus macaque white adipose tissue tend to be *cis-*regulatory elements for lipid metabolism-related genes, consistent with humans’ greater body fat percentage (Swain-Lenz et al. 2019). Such analysis of chromatin activity divergence has not been conducted on more recent evolutionary timescales within the human lineage, where mechanistic insights could aid understanding of ancestry-dependent disease prevalence (Reddy and Burakoff 2003; Krishnan and Hubert 2006; Ogdie et al. 2021).

Here, we use ATAC-seq and H3K27ac HiChIP, a combined measure of activity and contact frequency (Mumbach et al. 2016), to generate ABC scores linking candidate *cis-*regulatory elements (CREs) to candidate target genes (hereinafter “target genes”) in eight populations of African or European ancestry. We then decompose these scores into their activity and contact components to identify differential CREs (diff-CREs) for each score between individuals of African and European ancestry (Fig. 1a). Intersecting our diff-CREs with bQTL reveals three transcription factors (NF-κB, JunD, and PU.1) whose binding sites show signs of lineage-specific selection for differences in binding between the African and European ancestry populations. Our findings illustrate the utility of ABC scores to identify previously unappreciated population-specific activity of CREs, their target genes, and potential mechanisms of gene regulation.

## Results

### Differential activity-by-contact between African and European ancestry populations

We previously performed ATAC-seq in lymphoblastoid cell lines (LCLs) from ten different global population sequenced by the 1000 Genomes Project (Tehranchi et al. 2019). This was carried out in a pooled study design, with each population represented by a single pool of ∼100 unrelated individuals. We selected the four African (ESN, GWD, LWK, and YRI) and four European (CEU, FIN, IBS, and TSI) ancestry (hereinafter AFR and EUR, respectively) populations for comparison to isolate the effects of any lineage-specific selection on gene regulatory elements that has occurred since the divergence of human populations native to these two continents. The AFR and EUR ancestries were represented by 418 and 413 individuals, respectively. We first identified a common set of CREs by (1) calling peaks on ATAC-seq data combined across the four population pools of each ancestry, then (2) resizing them to 500 bp centered on each peak summit to avoid any potential peak width bias, and (3) retaining the top 150,000 by read count ranking. We ensured equal peak contributions between ancestries (see Methods) to balance statistical power and for consistency with how the ABC model was developed (Fulco et al. 2019).

To obtain the additional activity component and the contact component necessary for computing ABC scores, we performed H3K27ac HiChIP, which enriches first on the level of H3K27ac and second on HiC contact frequency of the two interacting regions (Mumbach et al. 2016), in two replicates per population of the same pooled LCLs (see Methods). We mapped reads from each replicate to a common reference. To minimize allelic mapping bias, we retained only reads overlapping variants that mapped to the same unique location after swapping out one allele for the other (van de Geijn et al. 2015). Subsequent filtering to reads in valid *cis* interaction pairs yielded ∼540 million paired-end reads qualified for use in ABC score computation (Fig. S1, Table S1).

To calculate ABC scores for each population, we jointly estimated activity level and contact frequency as the product of normalized ATAC-seq reads overlapping a given element and normalized HiChIP reads overlapping that element and the promoter of a given gene at 5 Kb resolution (see Methods). Of the 50,478 CRE-target gene (enhancer-gene) pairs with nonzero ATAC and HiChIP signal in all samples and passing enhancer-gene pair candidacy thresholds (see Methods) in at least one sample, 2,911 (∼5.8%) had differential ABC scores (diff-ABC) between AFR and EUR populations (diff-ABC P < 0.05, see Methods). Of these diff-ABC enhancer-gene pairs, 1,418 had higher ABC scores in AFR (AFR high ABC) and 1,493 had higher ABC in EUR (EUR high ABC). These AFR and EUR high ABC enhancer-gene pairs comprised 1,199 and 1,291 distinct target genes, and 1,122 and 1,184 distinct CREs, respectively.

### Differential CRE activity is linked to differential expression between ancestries

To determine the extent to which diff-ABC scores are associated with differential gene expression (DE) between African and European ancestry individuals, we analyzed gene expression data from two previous studies. Lea et al. (2022) measured gene expression across 12 cellular conditions (11 exposures and one unexposed control) in many of the same LCLs from African and European populations used in our study. Randolph et al. (2021) measured gene expression in non-infected (NI) and IAV-infected (flu) peripheral blood mononuclear cells (PBMCs) at single cell resolution from a panel of donors with varying degrees of African versus European ancestry. Both studies identified ancestry-associated DE (ancestry DE) genes, Lea et al. by modeling expression as a function of the African or European ancestry of each population, and Randolph et al. by modeling expression as a function of the proportion of African ancestry estimated from whole-genome sequencing. Although the context we assayed our LCLs in to generate ABC scores was closest to Lea et al.’s baseline/unexposed condition, by comparing diff-ABC in a baseline (unstimulated) context to DE in other contexts we were able to ask if CREs could be poised for DE regulation upon stimulation and/or in another cell type. We found our diff-ABC CRE target genes were enriched for ancestry DE in just five of the 22 contexts assayed across these studies (Fisher’s P = 2.10 x 10^-3^ - 6.79 x 10^-7^, Bonferroni-corrected P-value threshold = 2.27 x 10^-3^, see Methods; Fig. S2), none of which were the ABC-assayed context. These results suggest that diff-ABC CREs are generally not associated with DE due to the inclusion of activity level and contact frequency in ABC scores, absence of CRE poising, or some combination of the two.

ABC scores are designed to identify enhancer-gene pairs, but not the relative expression levels of target genes. We reasoned that decomposing each score into its components’ ATAC, H3K27ac ChIP, and HiC differential signal could increase concordance with ancestry DE by separating the contact component from the activity components to enable identification of cases where DE is linked to one of these mechanisms but not the other. Thus, for each enhancer-gene pair, we calculated separate scores using the ATAC-seq data (ATAC score), and using the HiChIP data to infer the H3K27ac level (ChIP score) and HiC contact frequency (HiC score) (Fig. 1a-b; see Methods). We then identified differential score (diff-score) enhancer-gene pairs as we did for diff-ABC (Fig. S3). To assess how each of these scores capture differences between populations and replicates we performed principal component analysis (PCA) and hierarchical clustering across samples on all enhancer-gene pairs for each score type. Since both CEU replicates were outliers (see Supplemental text, Fig. S4-7, Table S2), we removed this population, redefined enhancer-gene pairs, and computed scores for downstream analyses using the remaining 14 samples. PCA and hierarchical clustering on these scores shows ChIP scores are highly similar between replicates both when considering all enhancer-gene pairs (Fig. 1c) and the 5,000 most variable pairs, which also cluster by African and European ancestry. This clustering by ancestry holds when considering the 5,000 most variable enhancer CREs, but not promoter CREs (Fig. S8-9, Table S2-4). Although FIN rep1 and ESN rep1 are outliers for HiC scores, and thus also for ABC scores, we find little or no differential signal between ancestries among HiC scores both at the diff-HiC scores level (FDR = 0.87 at diff P < 0.05 relative to 0.093 and 0.057 for diff-ATAC and diff-ChIP, respectively; see Methods, Fig. S10, Table S5), and in downstream functional analyses.

We then tested if chromatin accessibility, H3K27ac levels, or HiC contact frequency had a stronger association with ancestry DE than diff-ABC did in the same 22 combinations of cell type and stimulation conditions. We found six enrichments for ancestry DE in diff-ATAC and five in diff-ChIP genes among the 22 tested contexts (P < 1.13 x 10^-3^, Fig. 2b). For example, target genes of diff-ChIP CREs were overrepresented among ancestry DE genes in LCLs after four hours of exposure to B-cell-activating factor (BAFF, odds ratio (OR) = 1.94, P = 6.1×10^-8^), a strong B cell activator and tumor necrosis factor family cytokine. For comparison, we also performed these enrichment tests for diff-ABC and HiC genes (Fig. S10). No contexts were enriched for ancestry DE in diff-HiC genes, and only DE genes in LCLs after four hours of exposure to ethanol (labelled “ETOH”) were enriched in diff-ABC genes at the same Bonferroni-corrected P-value threshold used for diff-ATAC and ChIP (Fig. S11, S12b). This indicates that the inclusion of the contact frequency component in ABC scores weakens the association of the activity components with DE. Overall, the strength of the associations of differential chromatin accessibility (diff-ATAC) and H3K27ac levels (diff-ChIP) with DE across several contexts suggests CREs could be poised for DE regulation upon stimulation or differentiation to another cell type.

**Figure 2.**
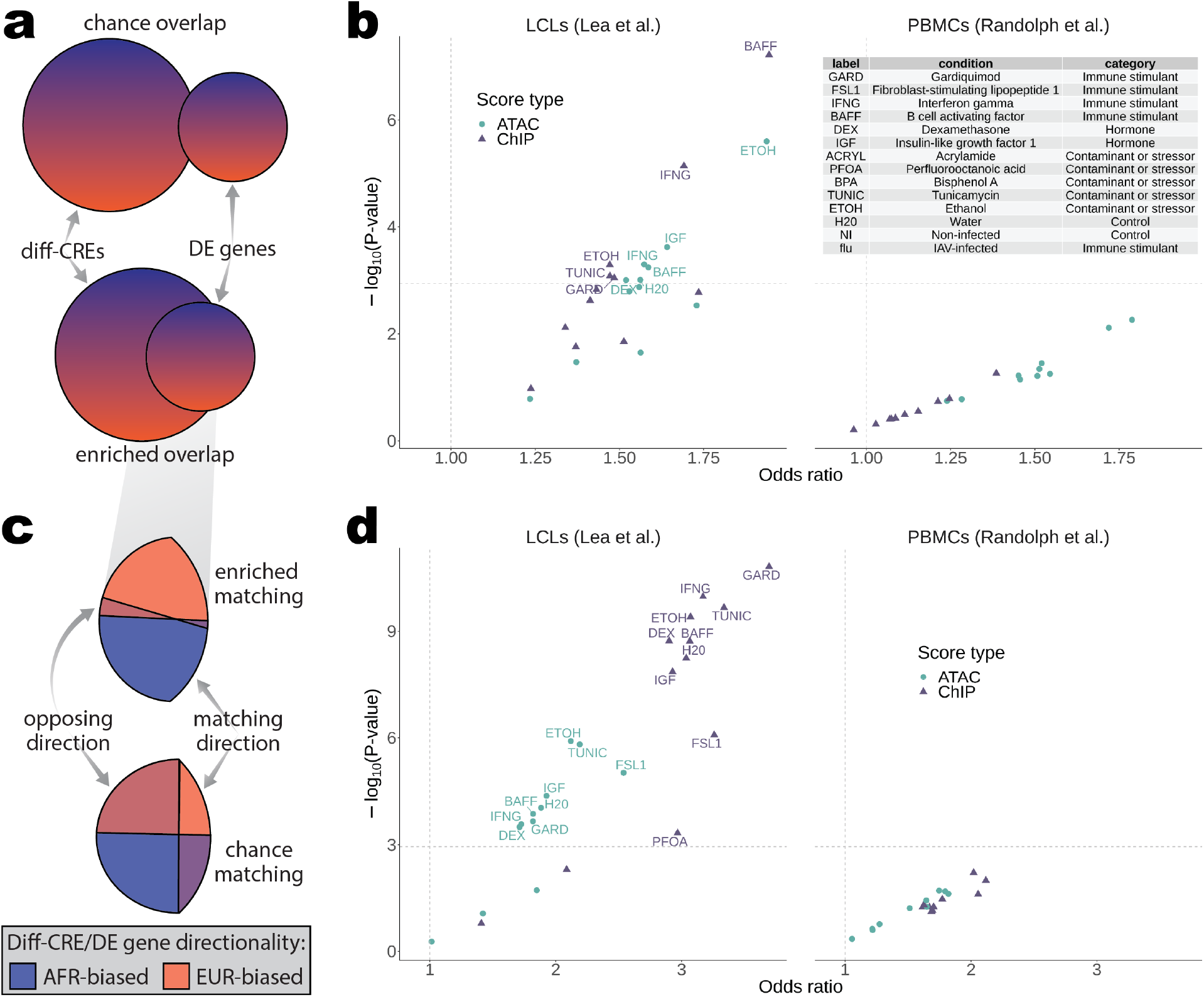
Differential enhancer and promoter enrichments for differential expression across conditions and cell types. **a)** Schematic depicting a simplified version of the enrichment analysis test results shown in (b). Any diff-CRE was counted as a “success” overlap in the hypergeometric enrichment test for a given context if it had a target gene that was DE in that context. Color gradients illustrate that each element has a directionality, which is the subject of analysis in (c) and (d). **b)** Results of one-sided Fisher’s exact tests for DE gene enrichment in diff-CREs are shown as odds ratios plotted against significance. Vertical dotted lines mark an odds ratio of 1 and horizontal dotted lines mark the Bonferroni-corrected P-value threshold. Points above this significance threshold are labeled with their context (see table inset). Score types used to defined diff-CREs in each test are indicated by the shape and color of the points. **c)** Schematic depicting the enrichment analysis test results shown in (d). Each element from any overlap in (a) (*e.g.*, gray gradient callout) was split into AFR- and EUR-biased directionality and any diff-CRE with a DE target gene in each context was counted as a “success” overlap if the ancestry DE direction matched the diff-score direction of the CRE (e.g., higher expression and higher score in AFR (blue)). For diff-CREs with multiple DE target genes, these targets were required to match direction to be included in each test. **d)** Results of one-side Fisher’s exact tests for diff-CRE matching DE directionality are shown as odds ratios plotted against significance. Vertical dotted lines mark an odds ratio of 1 and horizontal dotted lines mark the Bonferroni-corrected P-value threshold. Points above this significance threshold are labeled with their context. For a more detailed illustration of these tests, see Fig. S12a,c. differential CRE activity were driven in *cis* by any of these QTL types, as opposed to in *trans* by a difference in transcription factor expression level, we would expect strong associations between those QTL and differential activity CREs. We followed the same approach as in our DE analysis, first testing if our diff-CREs were enriched for any of these QTL relative to non-diff-CREs (Fig. 3a, see Methods). We found several significant bQTL enrichments across diff-ATAC and diff-ChIP CREs (Fig. 3b). We further asked if these enrichments were driven by bQTL in enhancers or promoters by performing separate tests on these two CRE types. Interestingly, enrichments for bQTL became even stronger when considering diff-ATAC and diff-ChIP enhancers, while diff-promoters showed no enrichments for any TF across all score types (Fig. 3c). These results suggest that many ancestry differences in CRE activity could be associated with differences in binding of specific TFs in *cis*, rather than differences in TF expression levels in *trans*.

Although gene expression can be predicted by promoter activity (Karlić, Chung et al. 2010), the contribution of promoter or enhancer activity to ancestry DE remains unknown. Thus, we asked if the associations between differential activity scores and differential expression were driven by genes whose top diff-CRE is a DE promoter. We found some evidence of this among diff-ATAC promoters across the non-infected and flu PBMC cell types (Randolph et al. 2021) (Fig. S13b), but no enrichments passed correction for multiple tests. We observed similar strengths of enrichment across the remaining contexts and score types for top diff-CRE enhancers and promoters (Fig. S13a-b); however, given only eleven significant enrichments when testing all CREs together we were likely underpowered to address this question.

Since the activity levels of enhancers and promoters usually increase and decrease with the expression levels of their target genes, we hypothesized that true enhancer-gene pairs would have higher expression in the same ancestry as that of the populations with higher ATAC and ChIP scores (Fig. 2c) and that this matching directionality would hold for pairs that are poised for DE in other contexts. To test this, we asked if among differential genes (diff-score FDRs = 0.093 and 0.057 for ATAC and ChIP, respectively; DE local false sign rate (LFSR) < 0.05) the ancestry direction of the top diff-CRE matched the DE direction of its target gene more often than expected by chance (see Methods). For example, is a gene with higher AFR ancestry expression also likely to have higher ATAC scores in AFR populations? We found that differential gene directionality matched more often than expected by chance in the same contexts in which differential activity and DE genes overlapped more often than expected by chance, as well as in three additional contexts for diff-ATAC and five for diff-ChIP. Six PMBC contexts (Randolph et al. 2021) were nominally enriched for diff-ChIP matching DE, though not significantly after multiple test correction. This was in contrast to genes identified by diff-ATAC CREs, which were only nominally enriched in five PBMC contexts (Randolph et al. 2021) at lower odds ratios (Fig. 2d, S12d). We found much weaker enrichment for matching DE directionality again among diff-ABC and HiC genes (Fig. S12d, S14).

To better ascertain the relative capacities of diff-ATAC and diff-ChIP (H3K27ac) to identify DE genes and their directionality, we compared the odds ratios across all contexts for all CREs and partitioned by promoter and enhancer top diff-CRE status. We found diff-ATAC to enrich better for DE (Wilcoxon P = 0.0022), whereas diff-ChIP enriched substantially better for matching DE direction (Wilcoxon P = 3.5 x 10^-6^) (Fig. S15, left). This difference held when considering enrichments derived from genes whose most differential CRE was a promoter but not an enhancer (Fig. S15, right and middle). While the diff-ChIP score incorporates H3K27ac levels from each target gene’s promoter for enhancer CREs (Fig. 1b, see Methods), diff-ATAC enhancers, which do not explicitly incorporate promoter accessibility information, performed similarly to diff-ChIP enhancers at identifying DE direction (Fig. S13c-d, S15). These results indicate that of the two types of chromatin activity assayed in our study, accessibility is the better indicator of which genes are DE, while H3K27ac levels better identify which ancestry has higher expression of these genes across numerous cellular contexts.

### Differential CRE activity is associated with ancestry-divergent variants that affect binding of specific TFs

Although diff-ATAC and diff-ChIP CREs are associated with DE of their target genes, the mechanism behind this association is unclear. To investigate this we sought to link potentially causal genetic variants to the activity of our CREs by intersecting them with QTL for the binding affinity of five transcription factors (bQTL, Fig. 3a) and H3K4me3 levels (H3K4me3 QTL) previously mapped in the same YRI LCLs used in our study (Tehranchi et al. 2016). If

**Figure 3.**
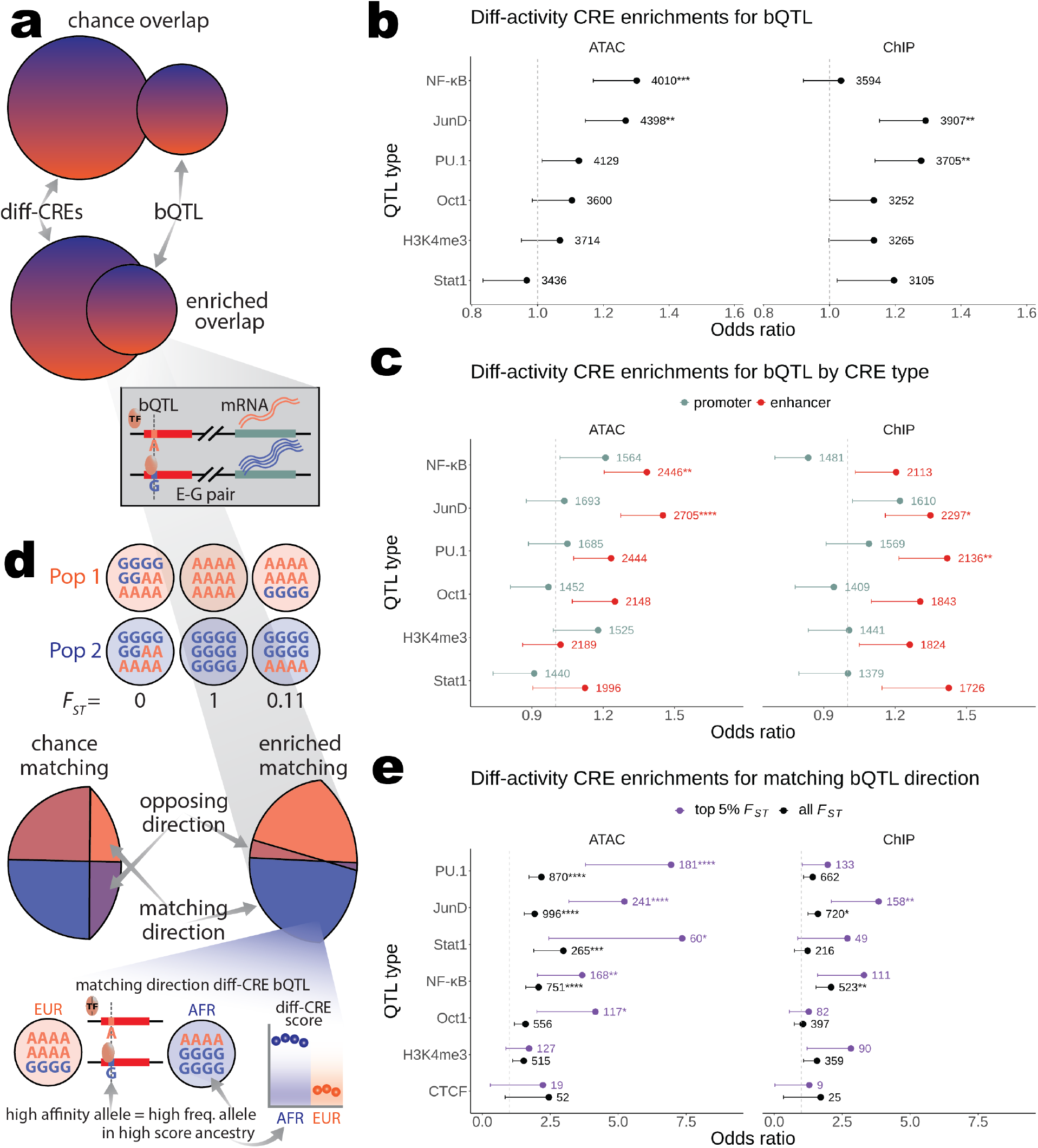
Enhancer differential activity is linked to sequence-dependent differences in NF-κB and JunD binding. **a)** Schematic depicting a simplified version of the enrichment test results shown in (b). Any diff-CRE was counted as a “success” overlap in hypergeometric enrichment tests if it contained a bQTL for the TF being tested (or H3K4me3 QTL). The boxed gradient callout from the enriched overlap shows a hypothetical example of a bQTL where the “G” allele increases the binding affinity of the TF, which leads to elevated mRNA levels through increased transcription of the target gene. **b)** TF bQTL and H3K4me3 QTL enrichments in diff-ATAC, ChIP, and HiC CREs are plotted as Fisher’s exact test odds ratios with error bars representing the lower bound of the 95% confidence interval. Since these are one-sided tests, the upper bound (infinity) is not shown. The total number of CREs used in each test is shown to the right of each odds ratio with asterisks indicating if the P-value passed multiple test correction (*, **, ***, and **** for Bonferroni-corrected P-value < 0.05, 0.005, 5×10^-4^, and 5×10^-5^, respectively). **c)** Results of tests on the same CREs used in (b) separated by their status as enhancer or promoter CREs. **d)** Schematic depicting the directionality matching enrichment test results shown in (e). Top, three hypothetical variants with “A” and “G” alleles in two populations where the first (leftmost) has no difference in allele frequency between populations or *F_ST_*=0, the second (middle) has the maximum possible difference in allele frequency between populations (“A” is fixed in population 1 and “G” is fixed in population 2) or *F_ST_*=1, and the third (rightmost) has the “G” allele at greater (but not fixed) frequency in population 2 or *F_ST_*≍0.11 in this case. Middle, diff-CRE bQTL from overlaps in (a-c) (gray gradient callout) were tested for if they match ancestry-associated direction more often than expected by chance. Bottom, a matching direction diff-CRE bQTL was defined as having its high affinity allele at higher frequency in the ancestry with the higher diff-CRE score. The blue gradient callout shows an example of a matching direction diff-CRE bQTL where the high affinity “G” allele from the same hypothetical bQTL shown in (a) is at higher frequency in AFR, which is the ancestry has the higher ATAC or ChIP scores defining the diff-CRE. **e)** Diff-CRE bQTL matching direction enrichment test results for diff-CREs defined by each displayed score type are plotted as Fisher’s exact test odds ratios with error bars representing the lower bound of the 95% confidence interval. Separate tests were performed on all diff-CRE bQTL (black) and those in the top 5% of *F_ST_* values among all variants in CREs genome-wide (purple). Since these are one-sided tests, the upper bound (infinity) is not shown. The total number of CREs used in each test is shown to the right of each odds ratio with asterisks indicating if the P-value passed multiple test correction (*, **, ***, and **** for Bonferroni-corrected P-value < 0.05, 0.005, 5×10^-4^, and 5×10^-5^, respectively).

To investigate the extent to which higher TF binding affinity corresponds to an increase in CRE activity, we asked if the high affinity bQTL allele was at higher frequency in the ancestry with higher CRE activity (Fig. 3d). We also included bQTL for CTCF (Ding et al. 2014), a protein that mediates chromosomal looping and chromatin, in these tests. The same TFs (JunD, NF-κB, and PU.1) were enriched for bQTL matching diff-CRE directionality as were enriched in diff-CREs overall, with the addition of STAT1 for matching diff-ATAC direction. Interestingly, PU.1 bQTL were enriched for matching diff-ATAC direction (Fig. 3e, left), but not diff-ChIP direction (Fig. 3e, right). This was in contrast with this TF’s overall bQTL enrichment in diff-ChIP CREs over non-diff (Fig. 3b-c), suggesting that this TF’s activity could be linked to context-dependent increases and decreases in H3K27ac levels, but is associated with increased chromatin accessibility in both cases.

If increased TF binding at bQTL is associated with an increase in CRE activity in *cis*, we should see an increase in correspondence between the ancestry with higher frequency of the high affinity bQTL allele and the ancestry with higher CRE activity the more extreme the difference in allele frequencies between ancestries. To test this, we asked if enrichments for matching directionality between bQTL and diff-CREs increase when considering only bQTL in the top 5% of *F_ST_* among variants in CREs (corresponding to *F_ST_* > 0.1813). Indeed, for all TFs with direction matching-enriched bQTL under no *F_ST_* thresholding we observed an average 2.36-fold increase in odds ratios when applying this *F_ST_* threshold (Fig. 3e). These enrichments were again driven by enhancers, as evidenced by the average 3.41-fold increase in odds ratios for the same comparison restricted to this CRE type (Fig. S16, top) and lack of directionality matching enrichment for any TF’s bQTL in promoters (Fig. S16, bottom). As expected, none of the above bQTL enrichment tests were significant for nominally diff-HiC CREs (Fig. S17).

Having established ancestry-dependent *cis* differences in TF binding as a likely mechanism for ancestry-associated differential CRE activity, specifically in enhancers, we sought to assess the likelihood that these TF bQTL have been under directional selection. We found that JunD, NF-κB, PU.1, and Oct1 bQTL have higher *F_ST_* in diff-ATAC than in non-diff-ATAC enhancers (Wilcoxon P = 2.4 x 10^-9^, 5.0 x 10^-4^, 2.2 x 10^-4^, and 6.0 x 10^-4^, respectively), consistent with differential binding of these TFs as drivers of differential enhancer activity. While none reached significance in diff-ChIP enhancers after multiple test correction, all of the QTL types except CTCF bQTL were nominally significant (Fig. 4a). This is likely due in part to power reduction in the diff-ChIP test due to the combination of lower resolution of HiChIP *cis* interaction pairs relative to ATAC-seq peaks (see Methods) and only counting the most significant bQTL per diff-score CRE (see Methods). Notably, there were no significant differences between bQTL in diff-versus non-diff promoters after multiple test correction. To assess evidence for selection on bQTL in diff-enhancers over those in diff-promoters more directly we performed the same test within diff-CREs between enhancers and promoters. Nearly all QTL types had higher median *F_ST_* in diff-enhancers than in diff-promoters for ATAC and ChIP although none were significant after multiple test correction (Fig. 4b). Again as expected, there was no difference in *F_ST_* between nominally diff- and non-diff-CREs or diff-CRE enhancers and promoters defined by HiC scores (Fig. S18). Overall, these higher *F_ST_* values for select bQTL in diff-enhancers are consistent with selection on TF binding sites in our diff-ATAC and diff-ChIP CREs.

**Figure 4.**
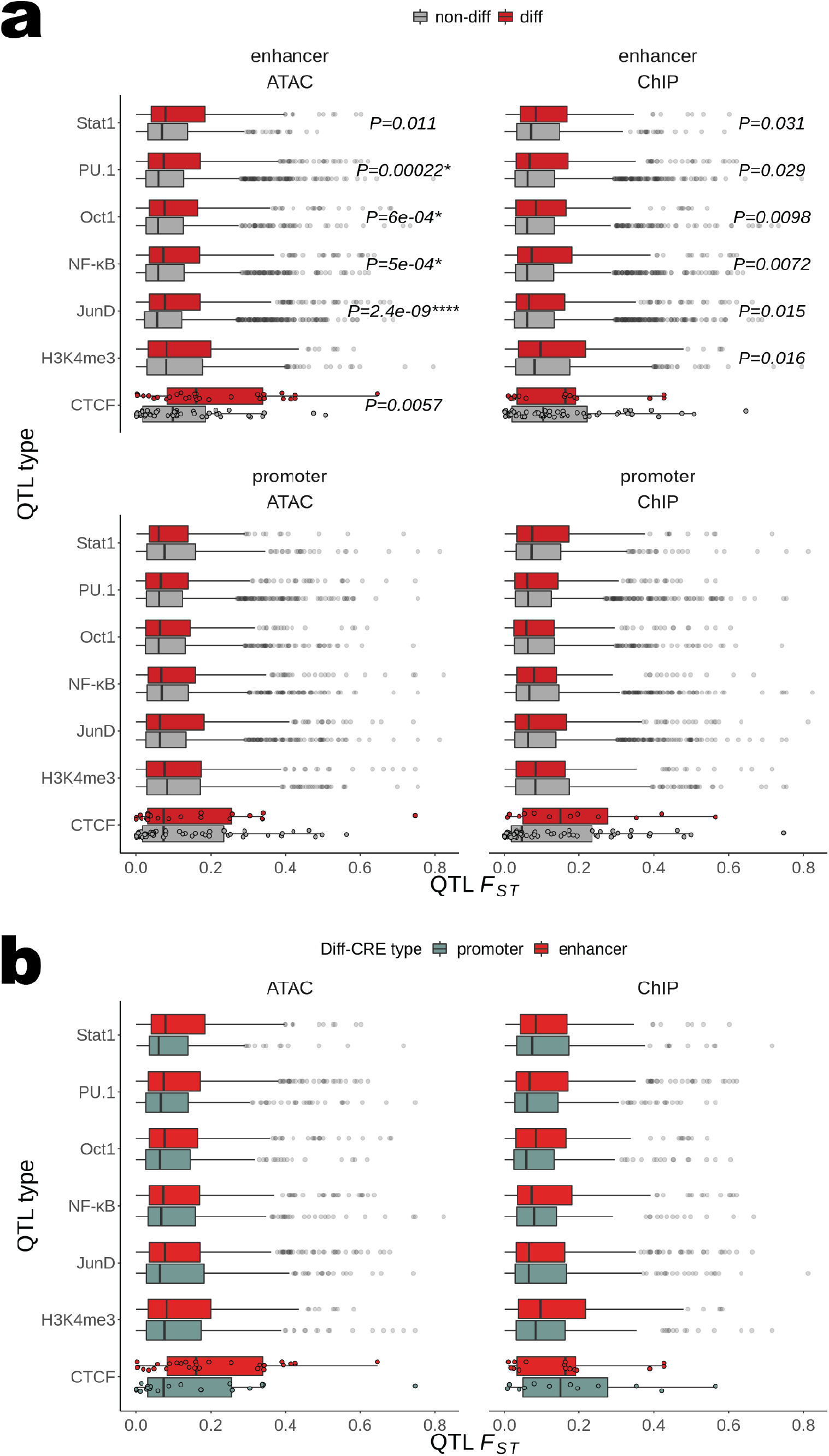
Evidence for selection on TF binding in enhancers versus promoters. **a)** *F_ST_* values of TF bQTL and H3K4me3 QTL are shown as boxplots for diff-and non-diff CREs of each displayed score type separated by if the QTL is in an enhancer (top) or promoter (bottom). The P-value from a one-sided Wilcoxon test on the *F_ST_* distributions from diff-versus non-diff CREs is displayed for all tests where diff-CRE bQTL *F_ST_* is nominally greater than that of non-diff (P < 0.05). **b)** *F_ST_* values of TF bQTL and H3K4me3 QTL are shown as boxplots for enhancer and promoter diff-CREs of each displayed score type. The P-value from a one-sided Wilcoxon test on the *F_ST_* distributions from diff-enhancer versus diff-promoter CREs is displayed for all tests where diff-enhancer bQTL *F_ST_* is nominally greater than that diff-promoters (P < 0.05). None of the P-values pass multiple test correction, so no asterisks are displayed. For visualization, non-outlier *F_ST_* values are only plotted over boxplots for distributions with fewer than 200 points.

### Differential CRE activity could be a result of directional selection and/or genetic drift

While these results could reflect directional selection, the underlying divergence in allele frequencies and corresponding ancestry-associated differential CRE activity could still be explained by genetic drift. More convincing evidence of directional selection could result from applying the sign test framework (Fraser et al. 2010; Fraser 2011) to ask if the high affinity alleles for bQTL that match diff-CRE direction are at higher frequency in one ancestry over the other more often than expected by chance. The sign test leverages the expectation that under neutrality, where genetic drift is the dominant force operating on allele frequency in populations of both ancestries, the high affinity alleles matching diff-CRE direction will not be biased toward higher frequencies in either population (Fig. S19a). Among bQTL matching diff-CRE direction, we found no more population-specific allele frequency bias than expected relative to the background of each TF’s bQTL in all CREs (Fig. S19b). Thus, genetic drift could be responsible for the association with increased ancestry divergence in diff-CREs matching bQTL directionality.

Moving from genotype up toward phenotype (Fig 1a, left), we next sought to identify the functional pathways most closely associated with our diff-score enhancer-gene pairs and their ancestry directionality. If variants in any subset of diff-CREs linked to target genes associated with a particular pathway have been subject to lineage-specific selection, these may not have been detected in our previous analyses. To address this possibility, we used gene set enrichment analysis with the gene ontology (GO) biological processes and MSigDB Hallmark gene sets on genes ranked by the difference in means between ancestries in ABC component scores of their top diff-CREs scaled by a measure of score variance (*i.e.*, ranked from high EUR activity to high AFR activity, see Methods). Again, under neutrality we would not expect diff-CREs with target genes in a particular pathway to have higher activity in one ancestry over the other. We did not find any significant enrichments among these gene sets after multiple test correction; however, some immune-related gene sets including interferon gamma (IFNG) response and TNF-*α* signaling via NF-*κ*B were among the top nominal enrichments for genes with top diff-ChIP and/or diff-ATAC CREs in the AFR high activity direction (Fig. S20). These nominal enrichments are consistent with diff-ATAC and diff-ChIP target gene enrichments for DE genes and matching DE directionality in IFNG-exposed LCLs (Fig. 2b,d, left), and Randolph et al.’s (2021) finding of TNF-*α* signaling via NF-*κ*B enrichment among genes with higher AFR expression in monocytes both before and after flu infection. Thus, although we do not find strong evidence for lineage-specific selection on diff-CREs in aggregate, the possibility that selection has more subtly affected gene regulatory architecture remains.

## Discussion

We have presented results from the first genome-wide comparison of chromatin activity and contact frequency between human populations with the goal of identifying CREs under recent selection. Since recent evidence points toward gene expression changes as the dominant force shaping recent human adaptation relative to protein sequence changes (Fraser 2013; Enard et al. 2014), this approach has the potential advantage of directly identifying CREs responsible for adaptive gene expression differences.

Using ABC scores to link CREs to target genes and decomposing these scores into their components allowed us to identify genes whose ancestry-associated expression differences across multiple contexts could be identified by the differential activity of their enhancers in the context of LCLs at baseline. This was particularly true for identifying the ancestry-associated direction of DE. Although H3K27ac alone is not required to maintain CRE activity (Zhang et al. 2020), it seems to be a more reliable indicator of expression direction than chromatin accessibility as measured by ATAC-seq in the context of our study. For example, one of many models capable of explaining this difference would be the binding of a transcriptional repressor to a promoter that yields an increase in chromatin accessibility but not in H3K27ac levels. About 25% of ABC-predicted and validated enhancer-gene pairs were found to have repressive effects via CRISPRi-flowFISH (Fulco et al. 2019) and any such effects within the matching DE directionality enrichments from our study could have contributed to the 39% of diff-activity pairs that “opposed” DE direction. More generally, the strength of these cross-context enrichments for DE and its direction is consistent with the maintenance of ancestry-associated regulatory differences in contexts beyond those where the target genes are DE. Matching differential CRE activity in LCLs at baseline and DE in many other contexts suggests CRE poising for DE regulation upon stimulation or differentiation to another cell type, or footprints of regulatory activity from a previous cell state remaining after the transition from that state.

Although our bQTL enrichment results suggest that differential activity is a result of cis-regulatory activity, it is possible that transcription factor differential expression in *trans* partially accounts for this. Indeed, JunD and NFKB2 (NF-κB subunit 2 of 2) show AFR-biased expression in LCLs at baseline (ancestry effect β = -0.26, LFSR = 0.0026 and ancestry effect β = -0.18, LFSR = 0.073, respectively); however, given the high odds ratios for bQTL in the top 5% of *F_ST_* (Fig. 3e, S16), differential CRE activity would likely persist even under constant *trans* conditions. Moreover, the lack of enrichments among diff-ATAC and diff-ChIP promoters for bQTL over non-diff (Fig. 3c), matching bQTL directionality irrespective of *F_ST_* (Fig. S16, bottom), and high bQTL *F_ST_* over non-diff (Fig. 4a), all relative to the positive enrichments found for tests on enhancers (including Fig. 4b) are consistent with greater evolutionary constraint on promoters and the distinct roles of enhancers in cell types that may be subject to different selection pressures (Heinz et al. 2015). While the ability of diff-ChIP enhancers to identify DE direction (Fig. S13c) is likely due to the inclusion of target gene promoters in ChIP scores, an analogous explanation cannot be made here since the bQTL in these enrichments are specifically in the enhancers, and not the target gene promoters. This genotype-level evidence restricted to differential enhancers indicates that our method of using chromatin as a spotlight on genetic variation effectively reveals otherwise hidden patterns consistent with selection (Fig 1a, left).

While our tests for greater transcription factor binding in one ancestry over the other did not show evidence of lineage-specific selection, the most enriched pathways among genes linked to higher activity CREs in AFR suggest more subtle effects of directional selection. For example, if the IFNG response pathway was under selection in one ancestry and this selection acted on a fraction of differential activity CREs regulated by transcription factor complexes more tissue- and/or response-specific than JunD or NF-*κ*B, this could remain undetected when aggregating over many more CREs. Our study is also limited by any changes to genome architecture introduced by Epstein-Barr virus in transforming B cells into LCLs that further mask the effects of any selection that has acted on B cells or even more relevant cell types and the noise introduced by combined analysis of multiple datasets generated by different people and/or labs. Future studies generating ABC score component data from diverse donors in cellular contexts more like those in which lineage-specific selection could have acted may find stronger evidence of it, especially if bQTL are mapped for more context-specific transcription factors.

The demographic processes that shape human genetic variation (*e.g.*, population history, migration, and drift) can obscure the influence of selection on variants that underlie adaptive phenotypes (Coop et al. 2009). Moreover, false signals of selection can result from under- controlled population stratification (Sohail et al. 2019; Berg et al. 2019). These confounders along with the prevalence of adaptive variants in non-coding regions with subtle effects (Fraser 2013) demonstrate the need for complementary methods to identify CREs that have been subjects of selection. We anticipate that extending the application of the method presented here to more populations and cell types will elucidate the molecular underpinnings of recent human evolution with implications for understanding modern disease prevalence.

## Conclusions

In generating the first population-level maps of candidate enhancer-target gene pairs in humans, we suggest *cis*-regulatory elements are poised for ancestry-dependent differential expression regulation upon stimulation or differentiation to another cell type. Mechanistically, this poising could be maintained by variants affecting the binding of transcription factors NF-κB, JunD, and PU.1 that show signs of lineage-specific selection in enhancers but not promoters. The potential effects of directional selection on immune-related pathways identified here suggest the promise of applying our chromatin-level selection test in additional cell types with roles in these pathways.

## Methods

### Cell culture and ATAC-seq

For detailed methods on cell culture conditions and processing, see our previous study (Tehranchi et al. 2019). Briefly, 2×10^3^ cells from each LCL were collected and pooled by population after growth to 6-8×10^5^ cells/mL. To prevent disproportionate cell line growth within pools throughout the collection and pooling process, sub-pools were frozen in liquid nitrogen at - 180 °C. After collection of all LCLs, sub-pools were combined by population and cells from each of the 10 pools were purified, isolated, and split into two replicates of 10^5^ cells each and pelleted according to Tehranchi et al. (2019) for a total of 20 samples. ATAC-seq was performed using the protocol from Buenrostro et al. (2015) in which each sample was resuspended in 100 ul of transposition mix containing 5 ul of Tn5 Transposase and incubated in a ThermoMixer for 30 min at 37 °C and 750 rpm. Transposed DNA fragments were then eluted and PCR-amplified with total cycles determined according to Buenrostro et al. (2015). Following two PCR cleanup steps, purified ATAC-seq libraries were sequenced on an Illumina HiSeq 4000 to generate 2×150 bp, paired-end reads.

### HiChIP

We thawed each −180°C-stored sub-pool described above and in Tehranchi et al. (2019) on ice, combined them by population and removed dead cells. As for ATAC-seq, to avoid disproportionate cell line growth we did not passage the cells before or after combining sub-pools. We then split each population pool into 2 replicates for crosslinking and HiChIP. For more detailed HiChIP methods, see Mumbach et al. (2017). Briefly, cells from each pool were pelleted and resuspended in 1% formaldehyde (Thermo Fisher) for crosslinking at a volume of 1 ml per million cells with incubation at room temperature for 10 min with rotation. Formaldehyde was then quenched with glycine at a 125 mM final concentration with 5 min room temperature incubation with rotation. Cells were then pelleted, PBS-washed, re-pelleted, and either used immediately in the HiChIP protocol or stored at −80 °C for HiChIP later.

HiChIP was performed as described in Mumbach et al. (2016) with H3K27ac antibody (Abcam, ab4729) and the following modifications. We used a 2 min sonication time, 2 µg of antibody, 34 µl of Protein A beads (Thermo Fisher) for chromatin-antibody complex capture. Post-ChIP Qubit quantification was performed to determine the amount of Tn5 used and number of PCR cycles performed for library generation, accounting for varying amounts of starting material. We performed size selection by PAGE purification (300-700 bp) to remove primer contamination and sequenced all libraries on an Illumina HiSeq 4000.

### ATAC-seq read mapping

For the complete mapping pipeline see https://github.com/kadepettie/popABC/tree/master/mapping, which contains a nextflow implementation of the steps described in Tehranchi et al. (2019). Cutadapt was used to remove sequencing adapters (arguments: -e 0.20 -a CTGTCTCTTATACACATCT -A CTGTCTCTTATACACATCT -m 5). PCR duplicate reads generated during library preparation were removed using picard MarkDuplicates (v2.18.20) (http://broadinstitute.github.io/picard/) (arguments: SORTING_COLLECTION_SIZE_RATIO=.05 MAX_FILE_HANDLES_FOR_READ_ENDS_MAP=1000 MAX_RECORDS_IN_RAM=2500000 OPTICAL_DUPLICATE_PIXEL_DISTANCE=100 REMOVE_DUPLICATES=true DUPLICATE_SCORING_STRATEGY=RANDOM). To minimize allelic mapping bias, a modified version (https://github.com/TheFraserLab/WASP/tree/atac-seq-analysis/mapping) of the WASP pipeline (van de Geijn et al. 2015) was used for read mapping. Reads were aligned to the hg19 genome using bowtie2 (Langmead and Salzberg 2012) (arguments: -N 1 -L 20 -X 2000 --end-to-end --np 0 --n-ceil L,0,0.15) and filtered to a minimum mapping quality of 5 using samtools (v1.8) (Danecek et al. 2021).

### HiChIP read mapping

HiChIP reads were mapped using the nf-core (Ewels et al. 2020) HiC-Pro (Servant et al. 2015) mapping pipeline (https://github.com/nf-core/hic) modified to include the same version of the WASP pipeline as was used for ATAC-seq to minimize allelic mapping bias (https://github.com/kadepettie/popABC/tree/master/hicpro). In this version however, allele-swapped remapping was performed separately on each read end, after which reads were re-paired, to accommodate the long-range nature of the paired-end reads as in the original HiC-Pro pipeline. Filtering reads down to valid *cis* interaction pairs, we took the raw 5 Kb resolution contact maps (the ‘.matrix’ and corresponding ‘.bed’ file output from process ‘build_contact_maps’) as input to our differential activity-by-contact pipeline.

### Differential activity-by-contact

#### Candidate element definition

We used the ABC model (https://github.com/broadinstitute/ABC-Enhancer-Gene-Prediction) to predict enhancer-gene connections in each pooled LCL population replicate (sample), with modifications to facilitate comparison of AFR and EUR population samples. For the complete differential activity-by-contact (diff-ABC) pipeline see https://github.com/kadepettie/popABC/tree/master/selection_1000G (“ABC_pipeline.nf”). We used Genrich (v0.5_dev, available at https://github.com/jsh58/Genrich) to call AFR and EUR ATAC-seq peaks jointly on the 8 samples from each with default parameters except for the following: -y -j -d 151. We then summed the reads in each peak across the corresponding 8 samples, kept the top 150,000 by read count, and resized them to 500 bp centered on the peak summit. To ensure equal contribution from peaks called separately in AFR and EUR to our candidate element input to the ABC model, we again made separate rankings by read count for each, then interleaved the two lists evenly by ranking, merging any overlaps and taking the top 150,000 elements. We next added 500 bp gene TSS-centered regions and removed any from the resulting list that overlapped regions of the genome with known signal artifacts (https://sites.google.com/site/anshulkundaje/projects/blacklists) (Amemiya et al. 2019; ENCODE Project Consortium 2012). Overlapping regions resulting from summit extensions and/or TSS additions were merged immediately following each of these steps. We defined promoter elements as those within 500 bp of an annotated TSS and the rest as enhancer elements.

#### Score component normalization

To ensure comparability of ABC scores between populations and replicates, and particularly samples with differing signal-to-noise ratios, we quantile normalized ATAC-seq reads per million sequenced reads (RPM), HiChIP valid *cis* interaction pair counts from 5 kb bins overlapping CREs, and each of these bins’ total count (for ChIP score computation) to the mean of their respective distributions across samples separately for enhancers and promoters. Since there is lack of consensus on an appropriate library size normalization method for HiChIP data, due to violation of the assumption of equal visibility between interacting regions often used in HiC normalization (Servant et al. 2015; Denker and de Laat 2016; Juric et al. 2019), we relied on the combination of quantile normalization and subsequent score normalization steps to control for library size and other technical artifacts.

Quantitative HiChIP signals were computed using the quantile normalized HiChIP contact counts according to Fulco et al. (2019). Briefly, for each gene TSS, all contact counts in CREs within 5 Mb were normalized to sum to one, then divided by the maximum of these values to normalize for comparison across genes.

#### Score computation

As in Fulco et al. (2019), we computed ABC scores using H3K27ac HiChIP with the fraction of regulatory input to gene *G* contributed by element *E* given by:

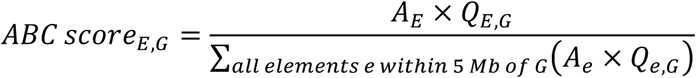

Here, the activity component (*A*) is quantile normalized ATAC-seq RPM, as in the original ABC score formula, but we have replaced the HiC contact component (*C*) with the quantitative HiChIP signal (*Q*) described above. We computed ATAC scores as follows:

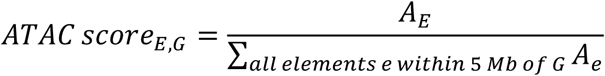

We computed ChIP scores as follows, using the geometric mean of the quantile normalized HiChIP bin totals overlapping each element (VC-sqrt) to estimate the aggregate H3K27ac signal at both elements:

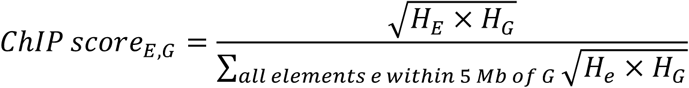

where *H* is the total quantile normalized valid *cis* interaction pair count from the HiChIP bin overlapping element *E* or the promoter of gene *G*. We computed HiC scores as follows, using HiChIP VC-sqrt normalization to estimate the contact frequency between each element independent of H3K27ac levels:

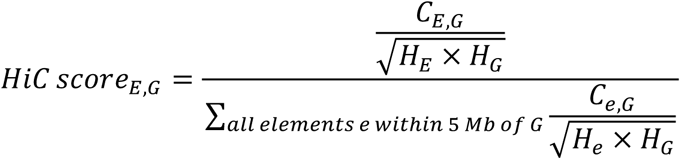

where *C* is the quantile normalized number of valid *cis* interaction pairs connecting the HiChIP bin overlapping element *E* and the promoter of gene *G*.

### E-G pair definition

To perform differential ABC score analysis across ancestries, we took predictions from the ABC model for each sample (population and replicate) and processed them according to the following steps. First, we excluded pairs with ABC score < 0.015 in all samples to avoid testing pairs unlikely to be true regulatory connections in any population (Nasser et al. 2021). Second, we excluded promoter-gene pairs with ABC scores below a stringent threshold of 0.1 because experimental data has shown the ABC model has poorer performance for this class of interactions, likely due to transcriptional interference, *trans* effects, and/or promoter competition (Fulco et al. 2019). Third, we required each enhancer-gene pair to be supported by non-zero quantile-normalized HiChIP contacts and ATAC values at the CRE in all samples, to avoid testing pairs where low ABC scores could be driven by mapping biases or low sequencing depth. Due to the difference in sequencing depths between samples, this final filtering step reduced the number of enhancer-gene pairs under consideration from 580,474 to our final set of 52,454 after removing CEU from the enhancer-gene pair-calling pipeline.

### Clustering analysis

For each score type and enhancer-gene pair, values were z-score normalized across samples for comparison and visualization of enhancer-gene pairs with large differences in mean score. PCA was performed with “prcomp” and heatmaps were generated using the *pheatmap* package (v1.0.12) in R (v4.1.0).

### Differential analysis

We called diff-CREs using unpaired, two-sample t-tests on each score type in AFR versus EUR samples. Log_2_ fold change effect sizes were estimated as the log_2_-ratio of the mean EUR score over the mean AFR score. We estimated a false discovery rate (FDR) for each score type at t-test P < 0.05 as the ratio of expected over observed enhancer-gene pairs with P < 0.05, where the null P-value distribution was derived from unpaired, two-sample t-tests on one set of replicates from each population versus the other. Replicate number was randomized for each enhancer-gene pair. To estimate the null P-value distribution for tests with eight AFR and six EUR samples after CEU removal while maintaining the eight versus six sample structure of each test, one AFR population was held out at random from replicate shuffling for each enhancer-gene pair and both replicates from this population were used in the group of eight (as opposed to the seven versus seven structure that would result from splitting by replicate across all populations). Since ChIP score signal is derived from HiChIP contact count bins at 5 Kb resolution, we counted diff-ChIP and HiC for CREs from the same HiChIP bin as one in each diff-score enrichment test described below.

### DE enrichments

We used hypergeometric tests (*i.e.,* one-sided Fisher’s exact tests) to determine enrichments for DE target genes among diff-CREs and matching ancestry directionality among DE genes with a diff-CRE. For the former across score types, we took the most differential CRE (top diff-CRE) by the corresponding metric (*i.e.*, ABC, ATAC, ChIP, or HiC score) per gene, defining diff-CREs at nominal t-test P-values < 0.05, non-diff-CREs at t-test P-values ≥ 0.5, and DE genes at LFSRs < 0.05 (Urbut et al. 2018; Randolph et al. 2021; Lea et al. 2022). Then, counting each CRE only once, we classified diff-CRE hits as any with at least one DE target gene, diff-CRE non-hits as any with no DE target genes, non-diff-CRE hits as any with no DE target genes, and non-diff-CRE non-hits as any with at least one DE target gene. For the promoter test, we took the subset of promoter CREs and additionally required diff-CRE hits and non-diff-CRE non-hits to be promoters for at least one of their DE target genes. For the enhancer test, we allowed promoter CREs to be classified as enhancers if they were not promoters for the relevant gene(s) (*e.g.*, a distal promoter for another gene contacting the promoter of the DE gene under consideration). That is, we required diff-CRE hits and non-diff-CRE non-hits not to be promoters for any of their DE target genes.

For the matching direction tests, we took the subset of top diff-CREs with DE target genes where all DE target genes were in the same direction (AFR- or EUR-biased) and, again counting each CRE only once, classified hits as diff-CREs with higher scores in the same ancestry as that with higher expression in their DE target gene(s). For the promoter and enhancer tests we required diff-CREs to be promoters for at least one of their DE target genes and none of their DE target genes, respectively. For each set of tests we only report P-values in the main text that pass Bonferroni-corrected thresholds.

### TF bQTL and H3K4me3 QTL enrichment analysis

We used hypergeometric tests to determine enrichments for each QTL type among diff-CREs relative to non-diff-CREs and matching ancestry directionality among diff-CREs with a QTL, using the same definitions for diff- and non-diff-CREs as in our DE enrichment analyses. In each test we considered the QTL with the lowest P-value per CRE for CREs with multiple QTL of the given type. For the directional analyses we defined bQTL directionality as AFR if the high affinity allele was present in AFR at greater frequency than in EUR and vice versa. For CREs with multiple bQTL, we additionally required that they all match direction for inclusion in each test. For binomial sign tests (Fig. S4a-b), we performed two-sided binomial tests on the number of QTL matching directionality in diff-CREs in the AFR direction out of the total number matching direction in diff-CREs with null probability of this proportion across all CREs.

### GO analysis

We used the R package fgsea (v1.20.0) (Korotkevich et al. 2021) to perform gene set enrichment analysis on genes ranked by the value of their most differential CRE according to the following statistic:

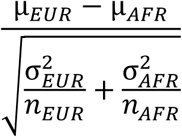

Where µ is the mean score, σ is the standard deviation, and *n* is the number of samples for each ancestry. fgsea was run on these ranked list for each score type using the C5 GO biological processes and MSigDB Hallmark gene sets with default arguments except: minSize = 15, maxSize = 500.

### *F_ST_* analysis

*F_ST_* for all variants was obtained using VCFtools’ calculation of Weir and Cockerham *F_ST_* (Weir and Cockerham 1984) between individuals from the African (ESN, GWD, LWK, and YRI) and European (CEU, FIN, IBS, and TSI) populations in our ATAC-seq and HiChIP data on a per-site basis. Variants with NA values were removed and negative estimations were adjusted to zero. For diff-versus non-diff CRE *F_ST_* Wilcoxon tests independent of their containing bQTL or H3K4me3 QTL, we took the maximum *F_ST_* value per CRE.

### Data and materials

- Genotype data for all individuals from populations used in our ATAC-seq and *F_ST_* analyses is from 1000 Genomes Project Phase 3 release (ftp://ftp.1000genomes.ebi.ac.uk/vol1/ftp/release/20130502/).
- All HiChIP reads are available as fastq files at NCBI SRA, project ID PRJNA898623 (https://www.ncbi.nlm.nih.gov/sra/PRJNA898623).
- Ancestry associated differential expression data from RNA-seq in LCLs after four-hour exposure to twelve cellular environments is from Lea et al. (2022) (https://genome.cshlp.org/content/suppl/2022/10/12/gr.276430.121.DC1/Supplemental_Table_S8.xlsx) with additional files and analysis code at https://github.com/AmandaJLea/LCLs_gene_exp.
- Ancestry associated differential expression data from single cell RNA-seq in PBMCs is from Randolph et al. (2021) (https://www.ncbi.nlm.nih.gov/geo/download/?acc=GSE162632&format=file).
- bQTL and H3K4me3 QTL are from Tehranchi et al. (2016) (https://www.cell.com/cms/10.1016/j.cell.2016.03.041/attachment/f7554f28-b82d-4b06-99b0-cbcfa7698d4c/mmc2.xlsx).
- CTCF QTL are from Ding et al. (2014).
- All pipelines and code used for analyses in this paper are available at https://github.com/kadepettie/popABC/tree/master.

## Acknowledgements

We thank members of the Fraser lab for helpful conversations, advice, and feedback on the manuscript; and Joseph Nasser, Kristy Mualim, and Jesse Engreitz for help with Activity-by-Contact scores.

## Contributions

HBF conceived the study. KPP and HBF conceived analysis methods. HiChIP experiments were performed by MM, MK, and KPP and funded by HYC. KPP performed all analyses and designed all graphics. AJL and JA provided unpublished data. KPP wrote the manuscript with input from all authors. HBF supervised all aspects of the work. All authors read and approved the final manuscript.

## Ethics declarations

### Ethics approval and consent to participate

Not applicable.

### Consent for publication

Not applicable.

## Supplementary information

### Supplemental text

#### Differential CRE activity is linked to differential expression between ancestries

##### Outlier removal

We observed that both CEU replicates were significant outliers in ChIP scores and to a lesser extent ATAC scores in PCAs and heatmaps of enhancer-gene pair scores (Fig. S4-7). CEU LCLs were the first established of any of the 1000 Genomes populations and their age has been proposed to drive their outlier status in previous studies (Yuan et al. 2015). Thus, to be conservative and ensure any subsequent results would not be driven solely by CEU, we removed this population, redefined enhancer-gene pairs, and computed scores for downstream analyses using the remaining 14 samples.

### Supplemental figures

**Figure S1.**
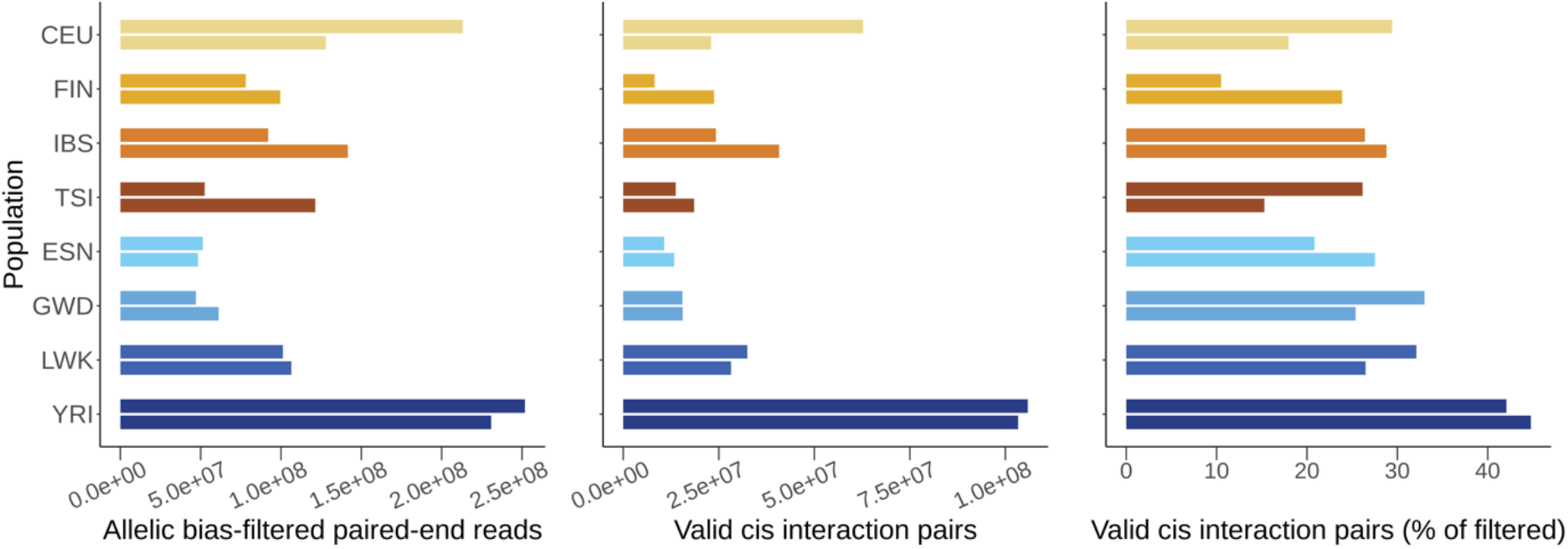
HiChIP reads filtering for use in ABC scores. Top bars are replicate 1 and bottom bars are replicate 2 for each population. Contact maps were generated from valid *cis* interaction pairs at 5 Kb resolution for computing ABC, ChIP, and HiC scores.

**Figure S2.**
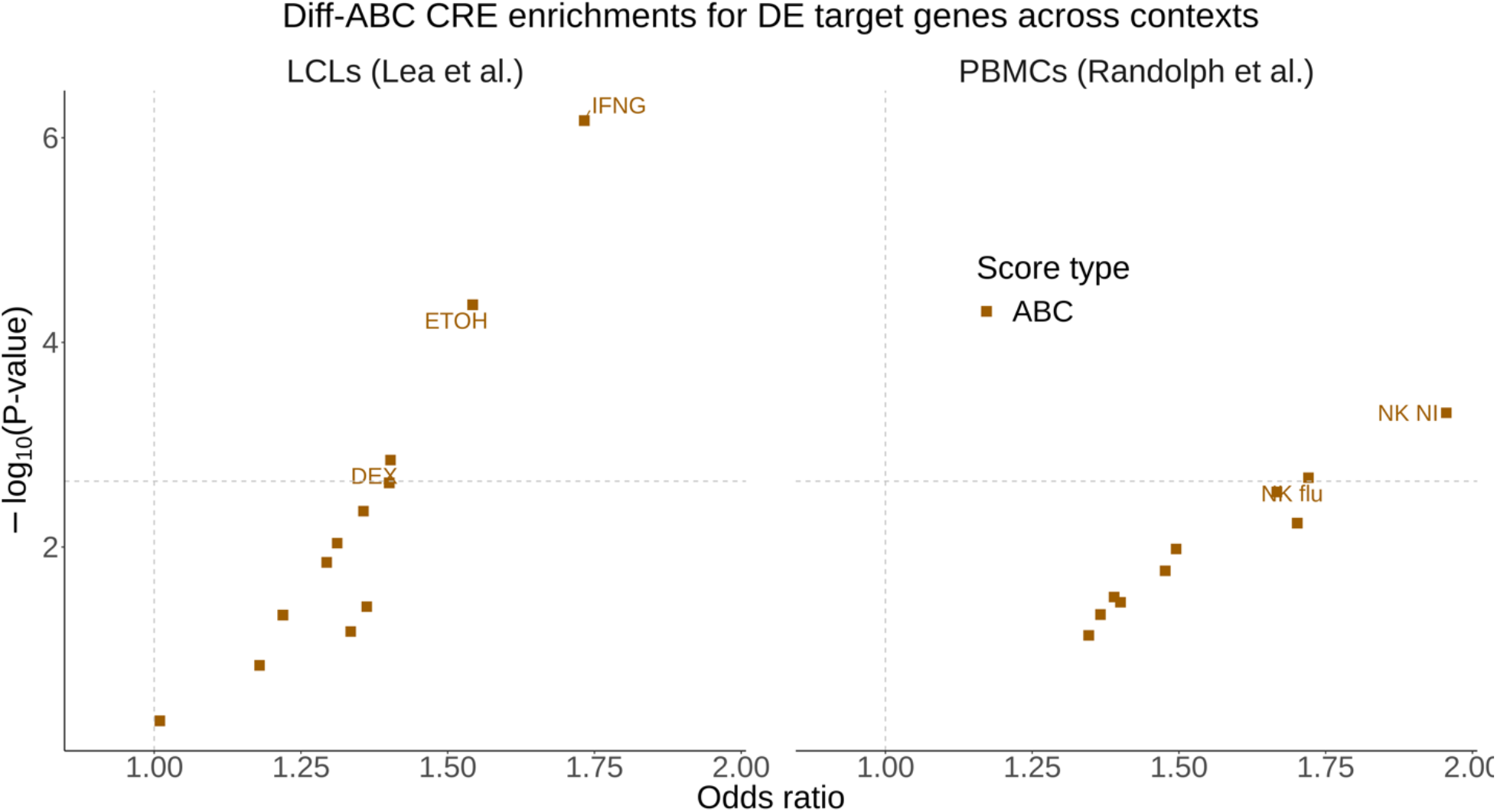
Diff-ABC (CEU-inclusive) enrichments for DE target genes across conditions and cell types. See Fig. 2a,b legend.

**Figure S3.**
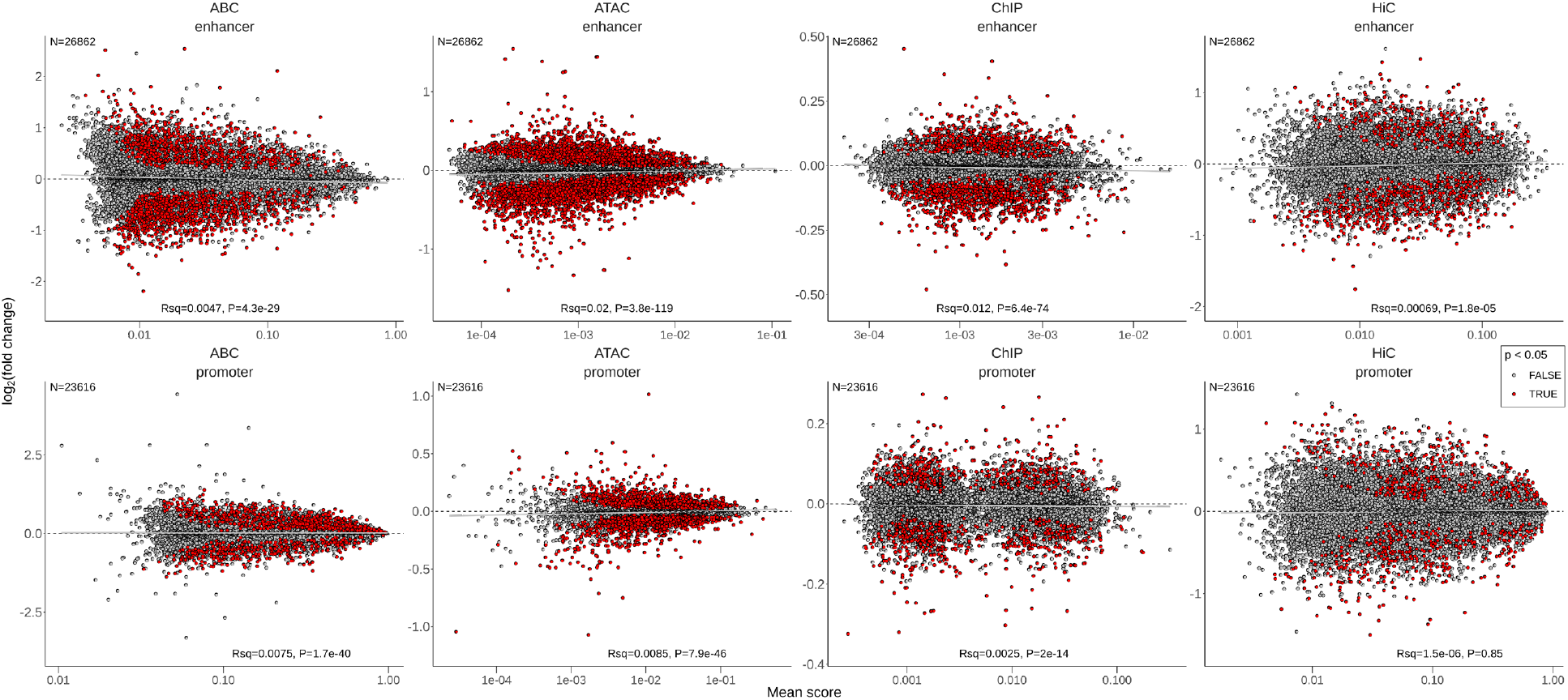
MA-style plots of diff-CRE (CEU-inclusive) directionality. The mean score (“A” scale) is plotted against the log_2_ ratio (“M” scale) or fold change of mean EUR score over mean AFR score for each CRE and score type. Red points are E-G pairs with diff-score P < 0.05.

**Figure S4.**
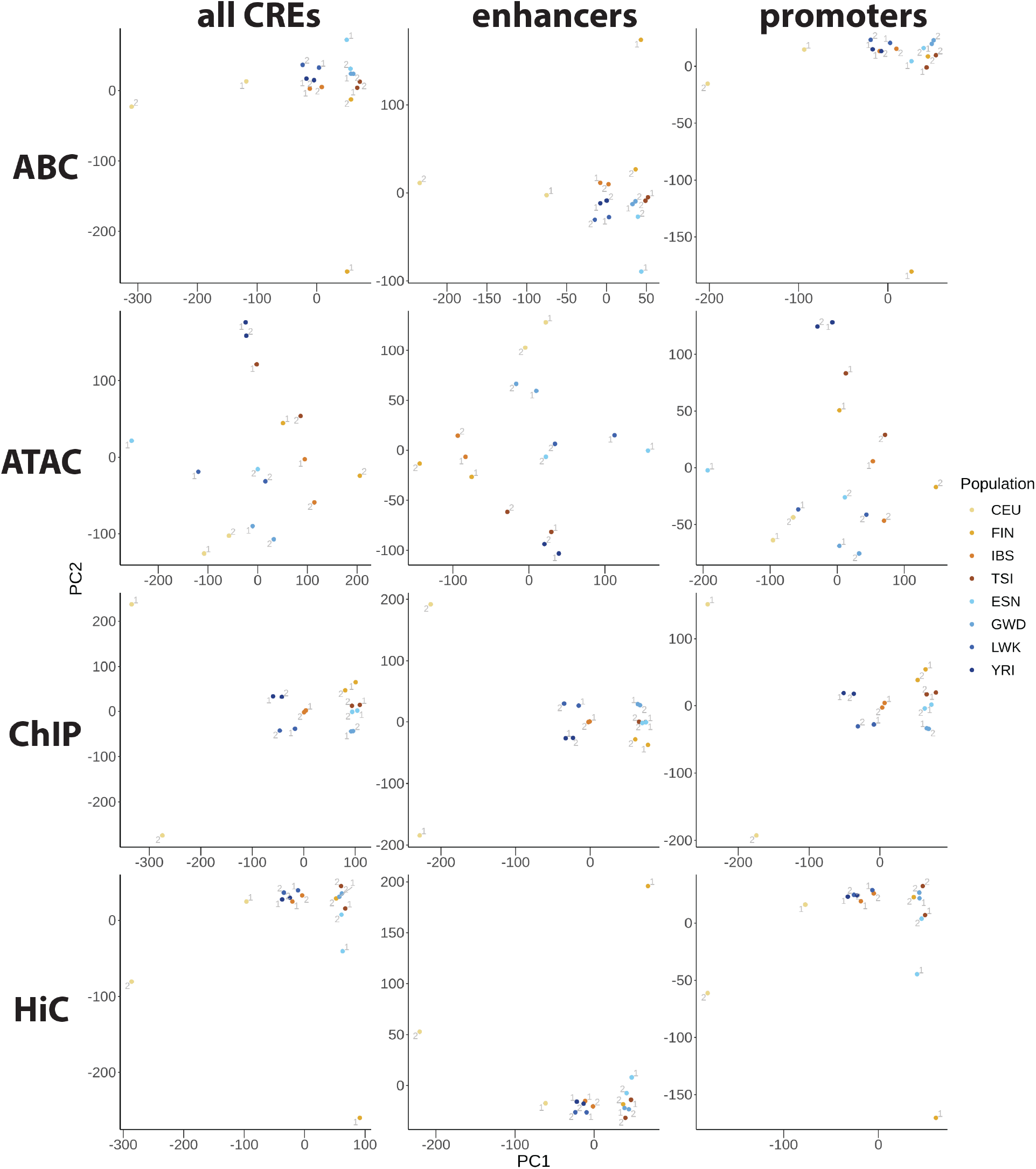
PCAs of all E-G pair scores (CEU-inclusive). Replicate numbers are displayed next to each point.

**Figure S5.**
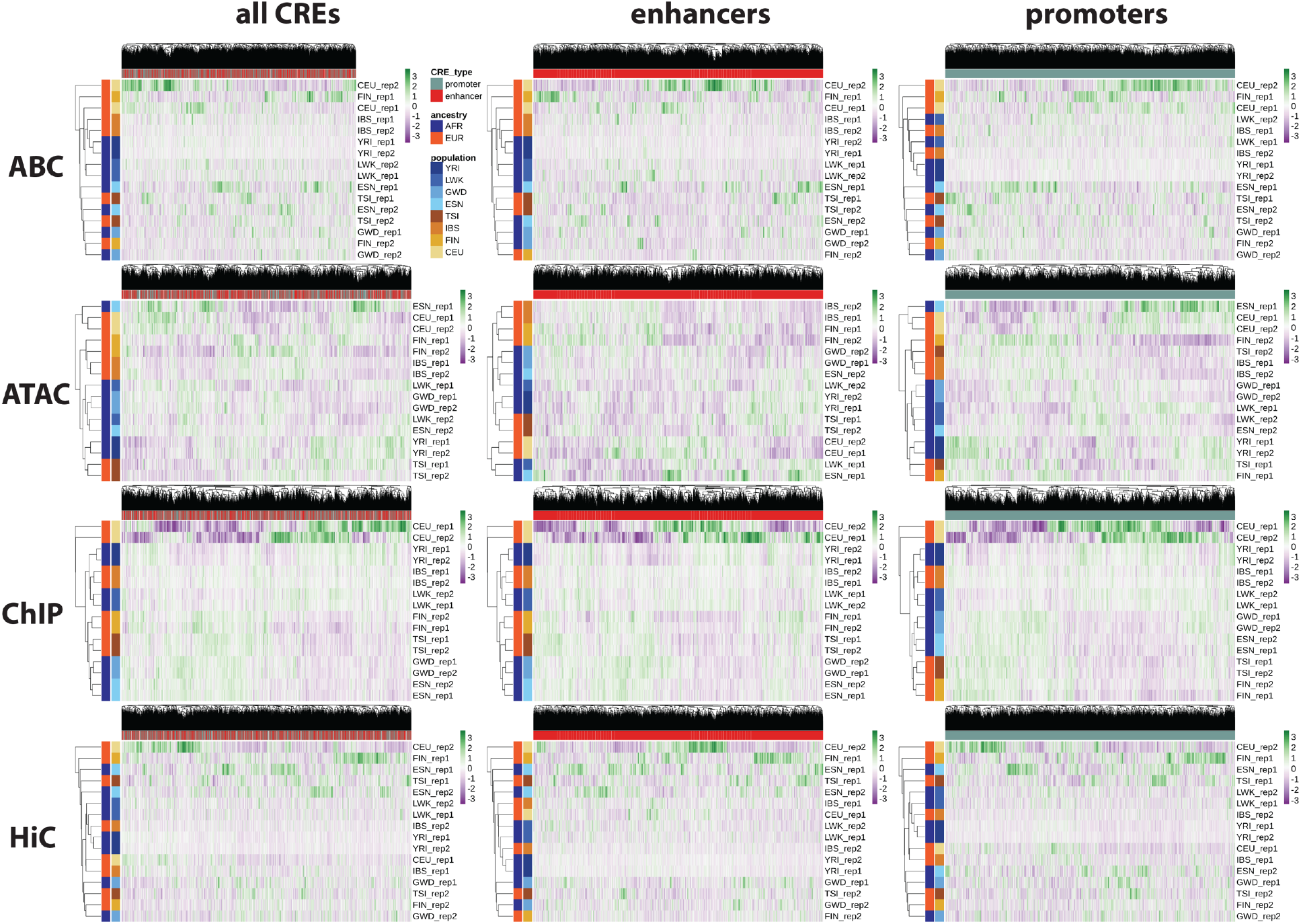
Heatmaps of all E-G pair scores (CEU-inclusive) with hierarchical clustering.

**Figure S6.**
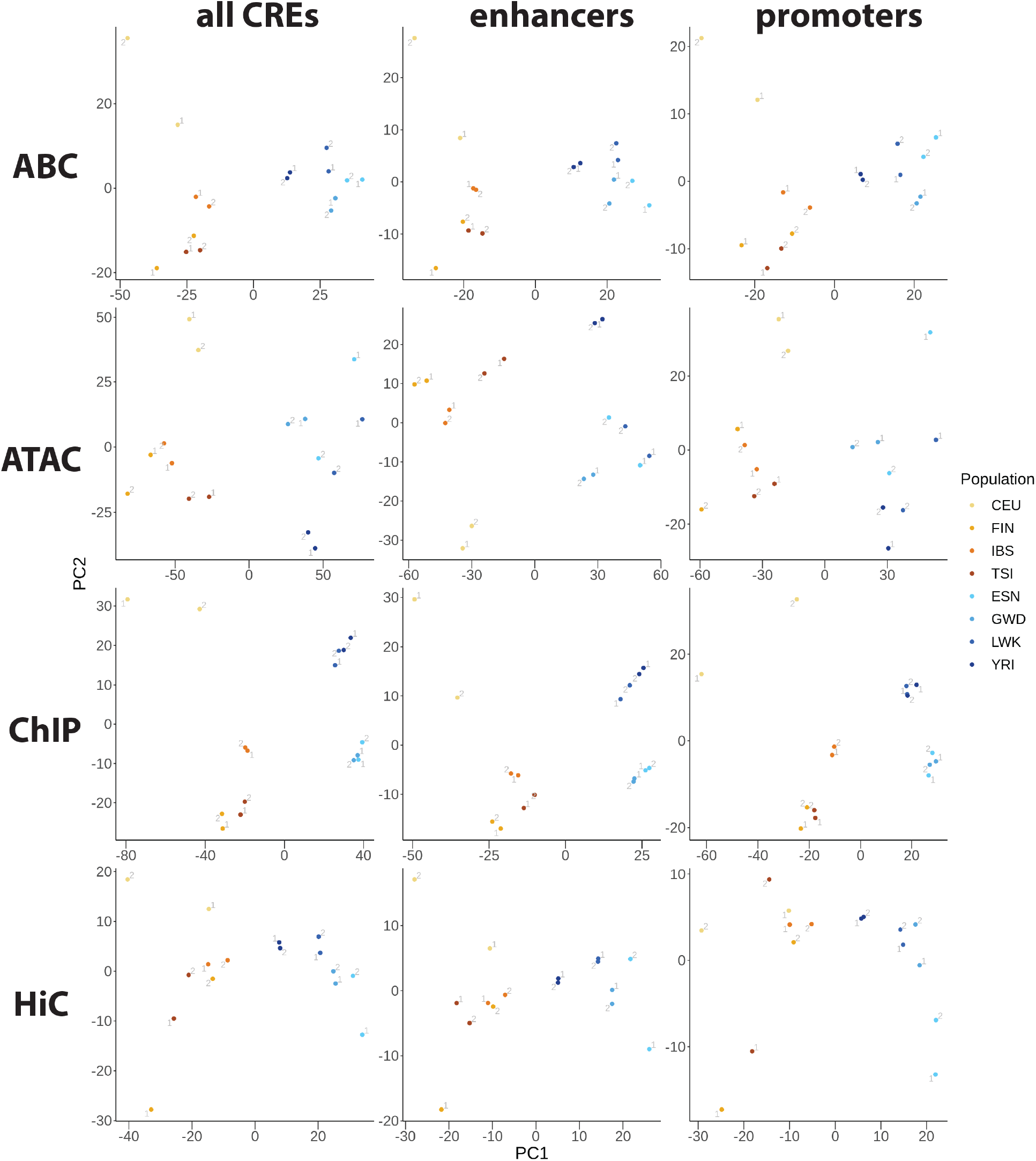
PCAs of top diff-E-G pair scores (CEU-inclusive). E-G pairs were subset first by diff-score P < 0.05, then by lowest P-value per gene, then by CRE type indicated by column names before performing PCA on each score type (row names). Replicate numbers are displayed next to each point.

**Figure S7.**
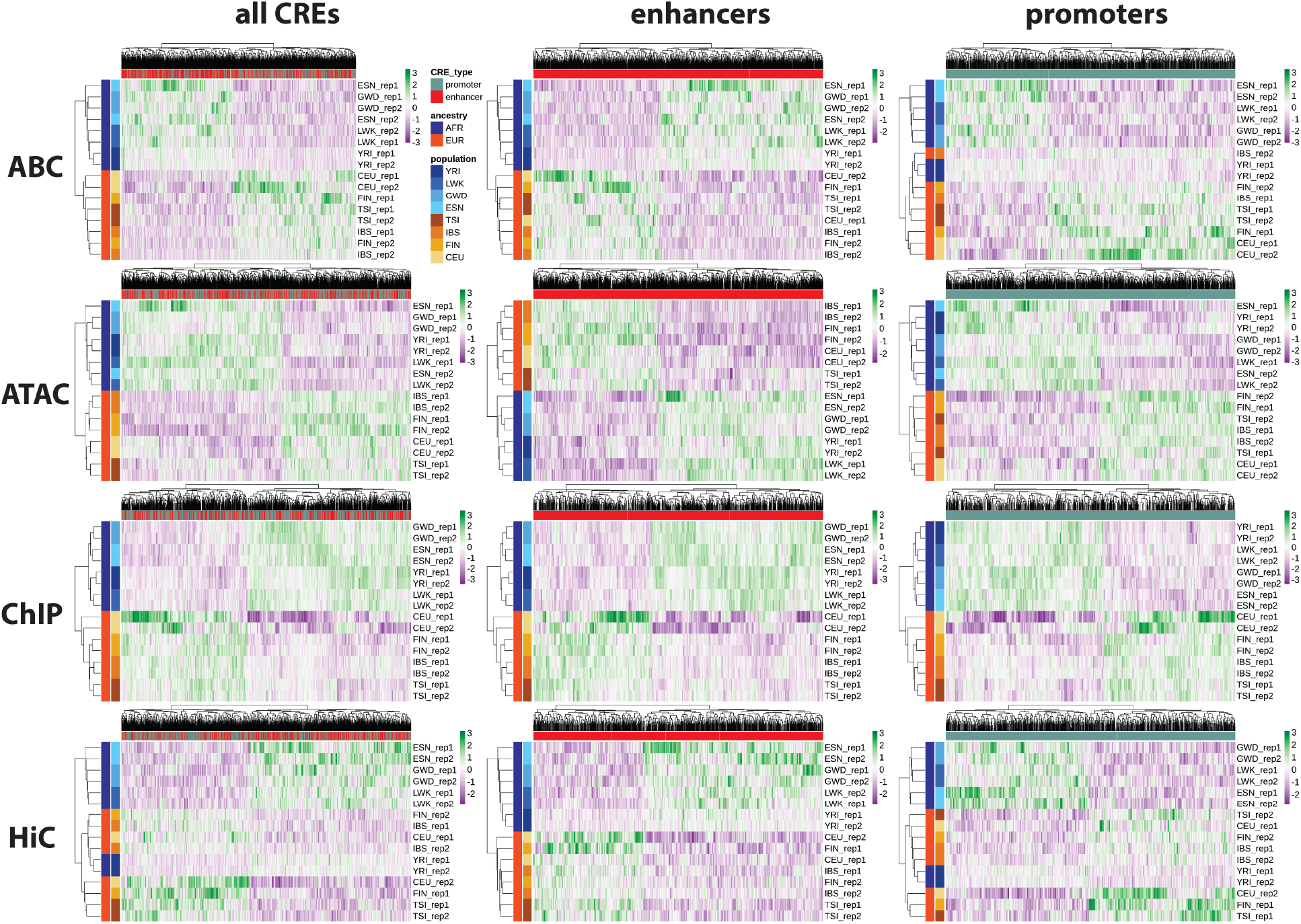
Heatmaps of top diff-E-G pair scores (CEU-inclusive) with hierarchical clustering. E-G pairs were subset first by diff-score P < 0.05, then by lowest P-value per gene, then by CRE type indicated by column names before performing PCA on each score type (row names).

**Figure S8.**
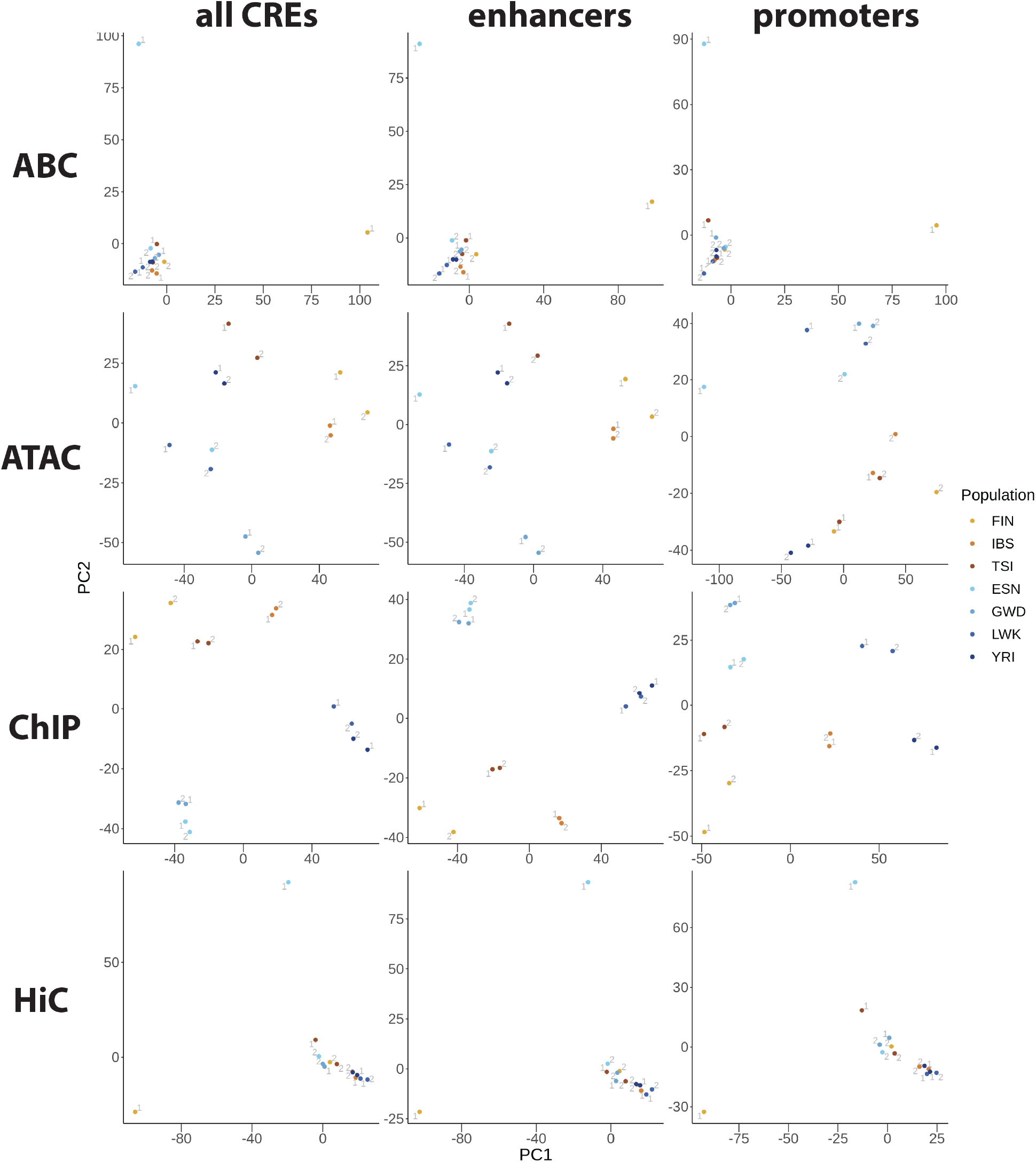
PCAs of the 5,000 highest coefficient of variation E-G pair scores of each CRE type. Replicate numbers are displayed next to each point.

**Figure S9.**
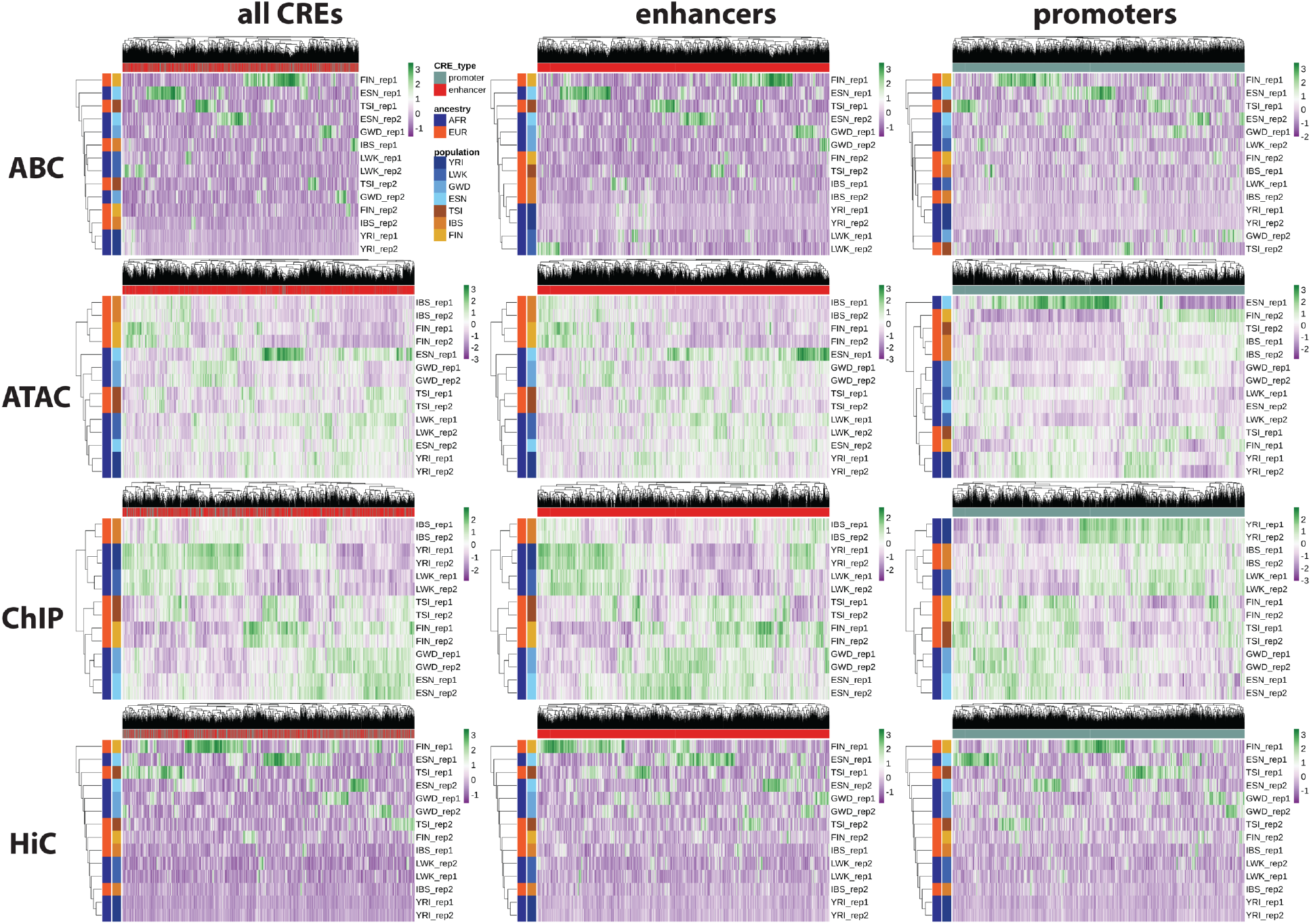
Heatmaps of the 5,000 highest coefficient of variation E-G pair scores of each CRE type with hierarchical clustering.

**Figure S10.**
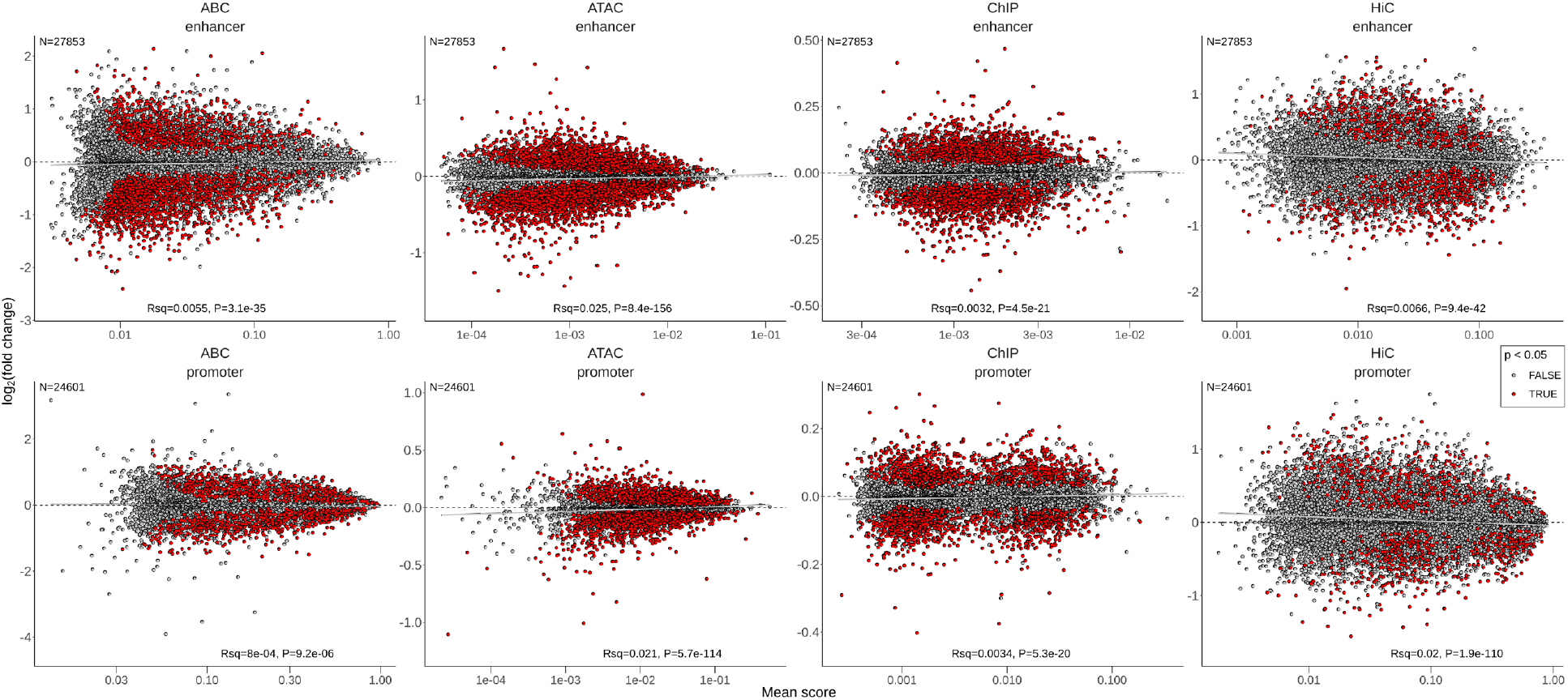
MA-style plots of diff-CRE directionality. The mean score (“A” scale) is plotted against the log_2_ ratio (“M” scale) or fold change of mean EUR score over mean AFR score for each CRE and score type. Red points are E-G pairs with diff-score P < 0.05.

**Figure S11.**
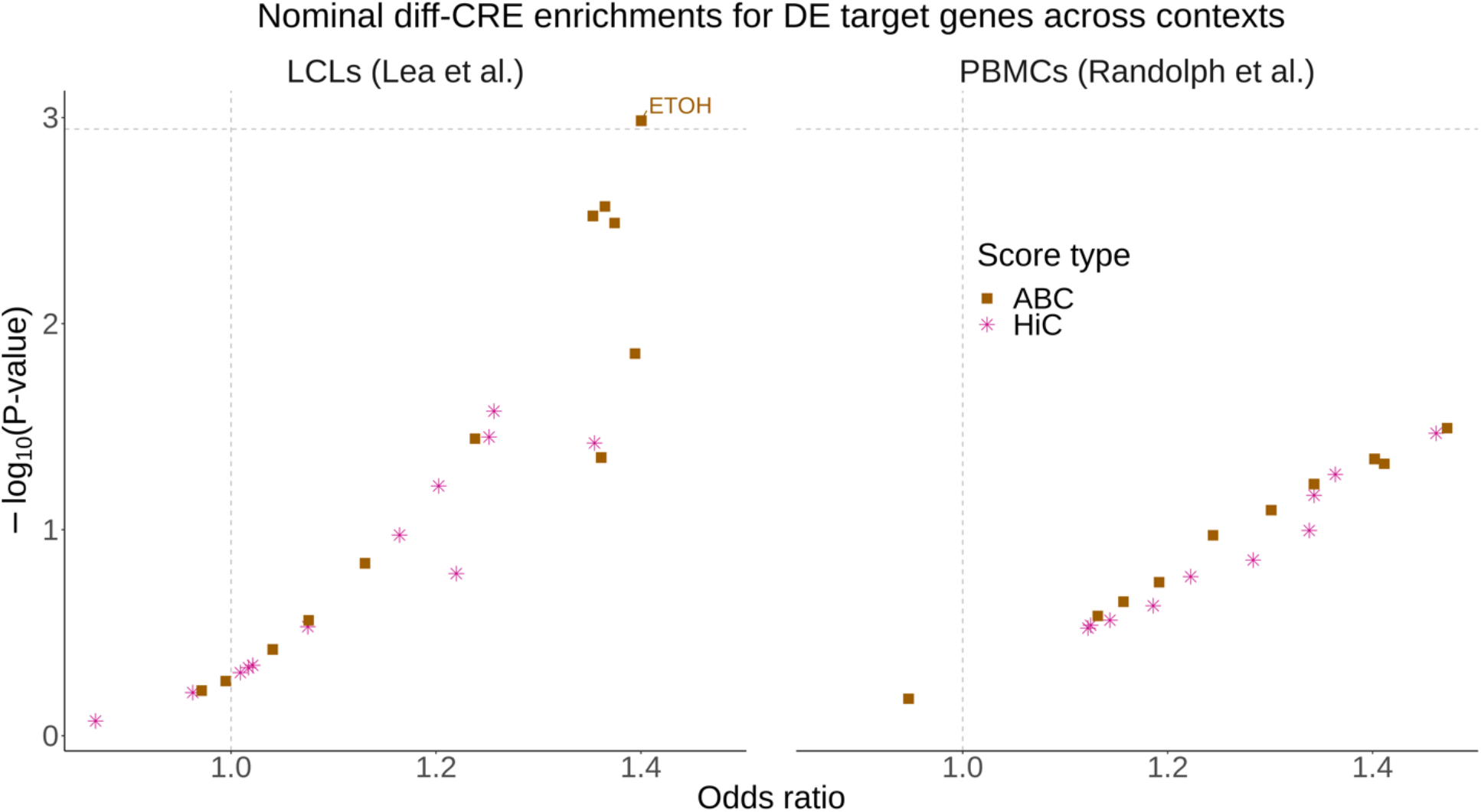
Nominal diff-ABC and HiC enrichments for DE target genes across conditions and cell types. See Fig. 2a,b legend. Horizontal dotted lines representing the Bonferroni corrected threshold are shown only for comparison and consistency with the main text figures, since these were not hypothesis tests due to the high diff-ABC and HiC FDRs (0.52 and 0.87, respectively, at diff P < 0.05).

**Figure S12.**
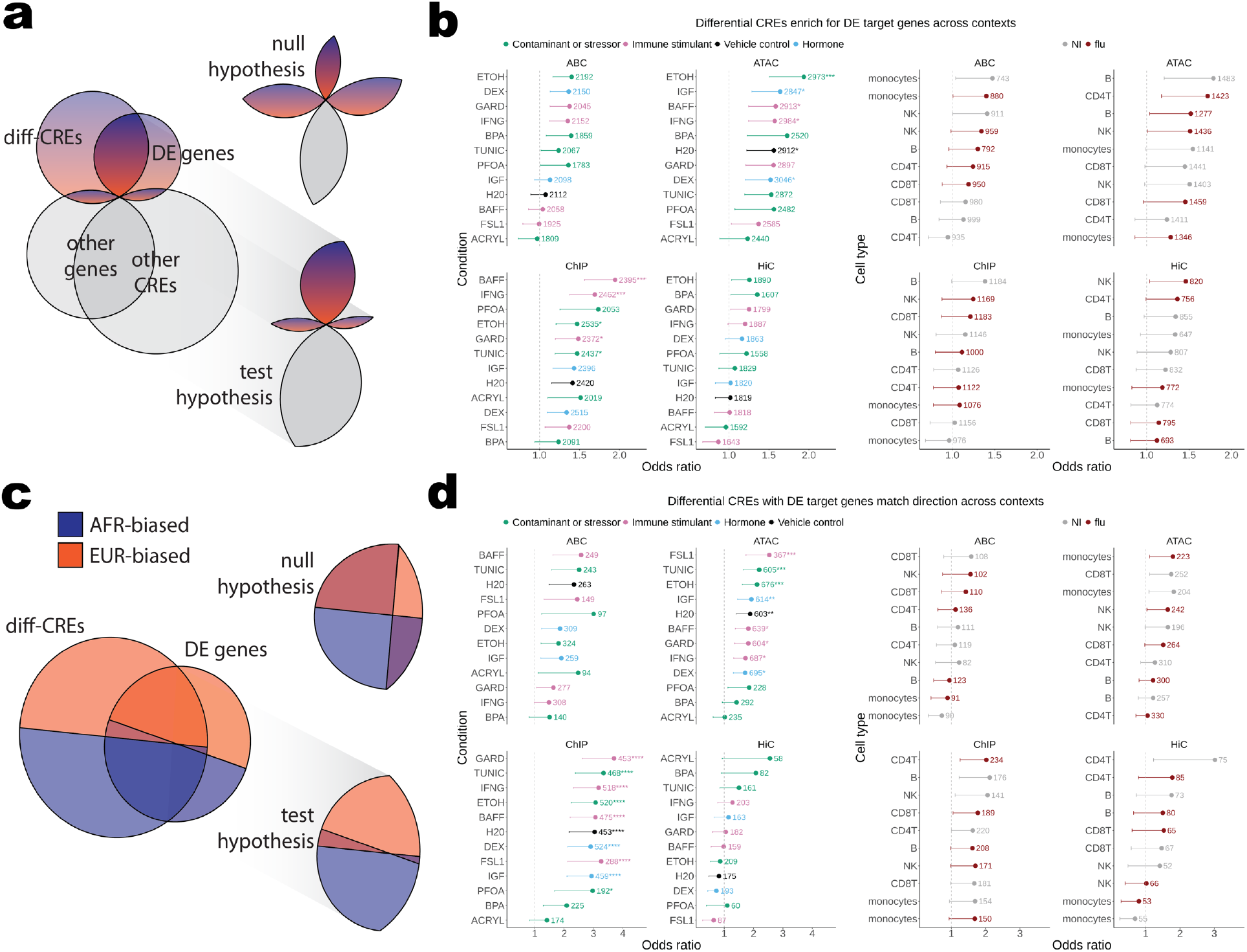
Diff-CRE enrichments for DE across conditions and cell types. See Fig. 2 legend, except here results of one-sided Fisher’s exact tests for DE gene enrichment in diff-CREs (b) and diff-CRE matching DE directionality (d) are plotted as odds ratios with error bars representing the lower bound of the 95% confidence interval. Since these are one-sided tests, the upper bound is infinity and is not shown. The total number of CREs used in each test is shown to the right of each odds ratio with asterisks indicating if the P-value passed multiple test correction (*, **, ***, and **** for Bonferroni-corrected P-value < 0.05, 0.005, 5×10^-4^, and 5×10^-5^, respectively). Nominal diff-ABC and HiC enrichments are shown for comparison only and are not included in Bonferroni correction, since these were not hypothesis tests due to the high diff-ABC and HiC FDRs (0.52 and 0.87, respectively, at diff P < 0.05).

**Figure S13.**
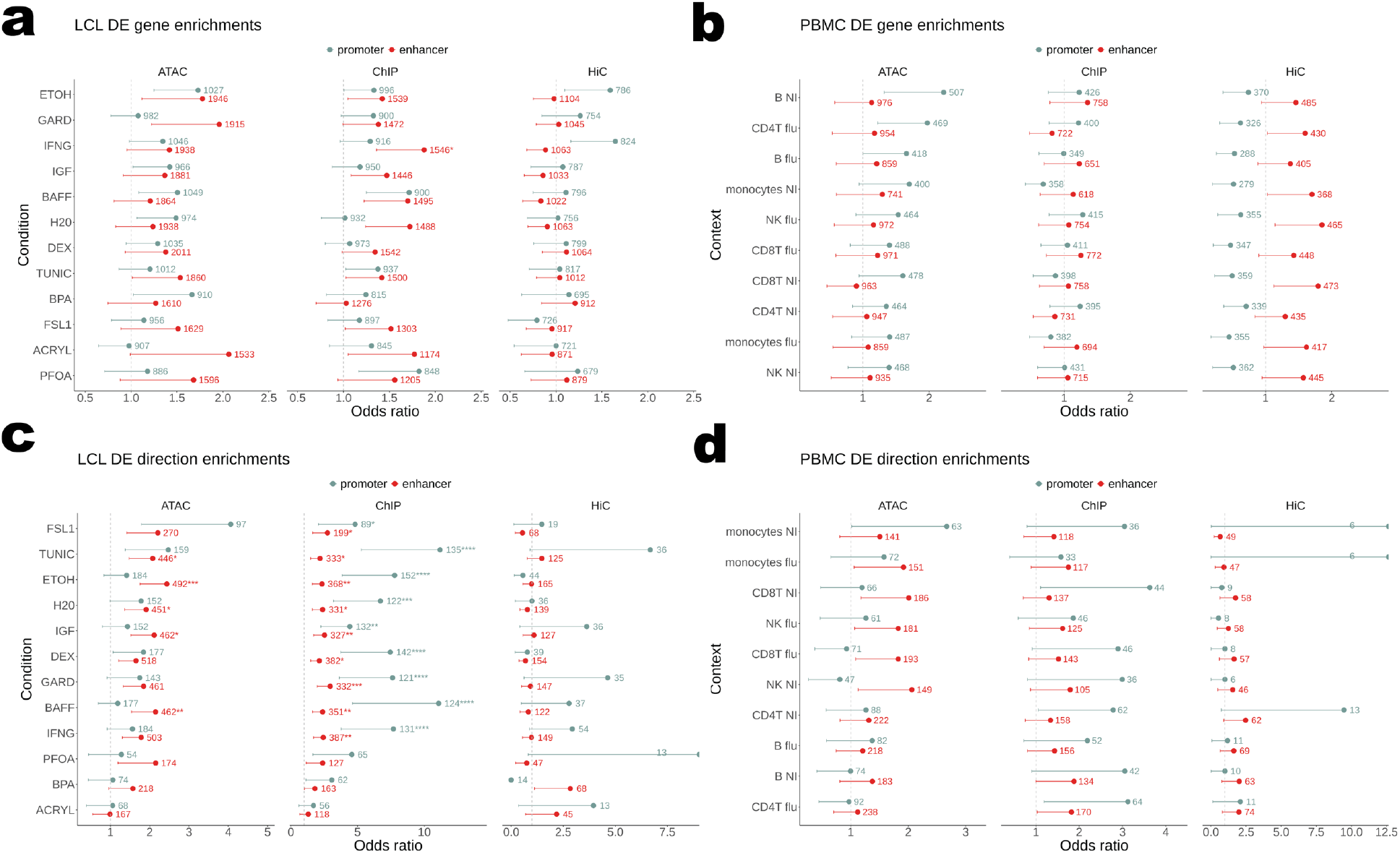
Diff-CRE enrichments for DE across conditions and cell types by CRE type. The same data used for tests in Fig. 2 and S12 were split into genes whose top diff-CRE was a promoter or enhancer and the same tests were performed on each, thus doubling the number of tests. See Fig. S12 legend. Nominal HiC enrichments are shown for comparison only and are not included in Bonferroni correction, since these were not hypothesis tests due to the high HiC FDR (0.87 at diff P < 0.05).

**Figure S14.**
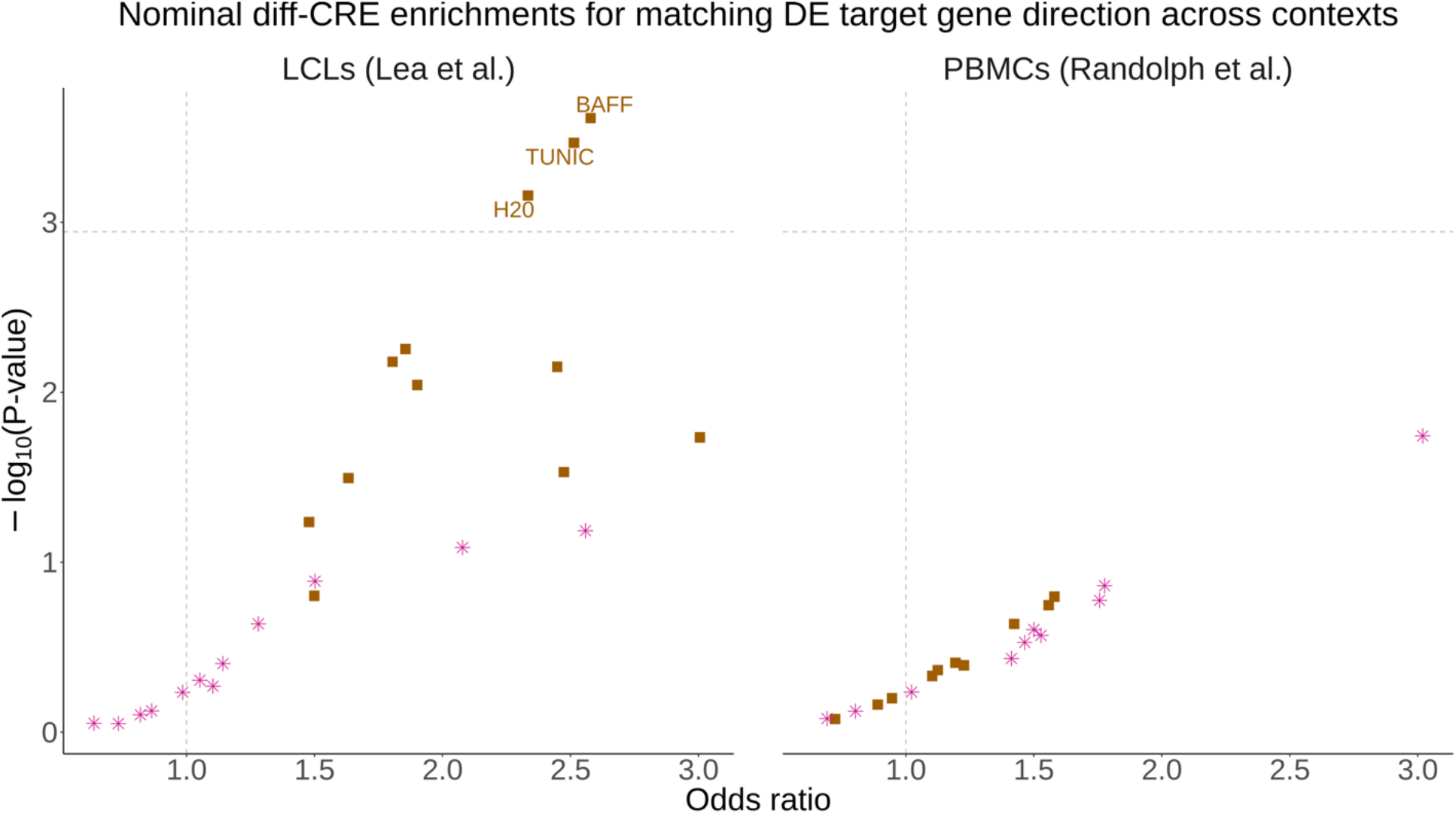
Nominal diff-ABC and HiC enrichments for matching DE target gene direction across conditions and cell types. See Fig. 2c,d legend. Horizontal dotted lines representing the Bonferroni corrected threshold are shown only for comparison and consistency with the main text figures, since these were not hypothesis tests due to the high diff-ABC and HiC FDRs (0.52 and 0.87, respectively, at diff P < 0.05).

**Figure S15.**
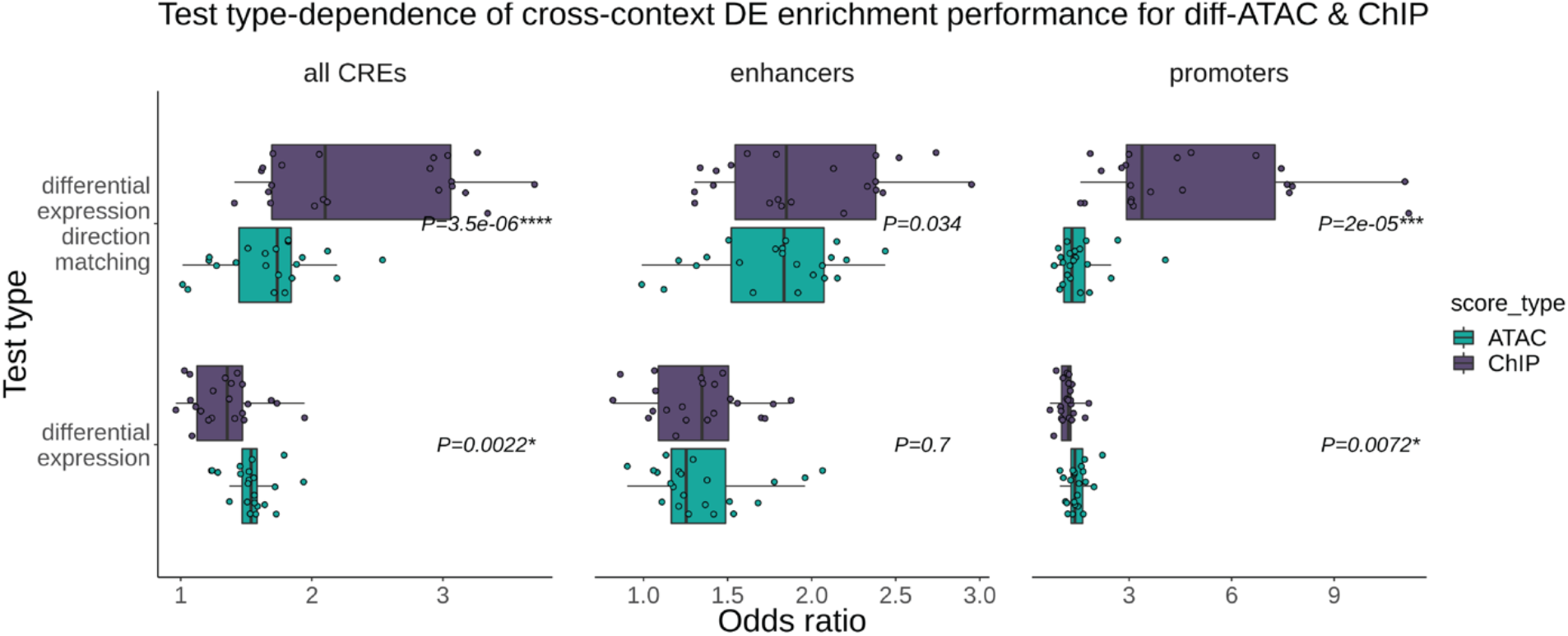
Comparison of odds ratios between diff-ATAC and diff-ChIP from DE gene and matching DE directionality enrichment tests. Odds ratios from tests in Fig. 2d are plotted as boxplots for each test type and CRE type. P-values from Wilcoxon tests on diff-ATAC versus diff-ChIP odds ratios in each category are shown. Asterisks indicate if the P-value passed multiple test correction (*, **, ***, and **** for Bonferroni-corrected P-value < 0.05, 0.005, 5×10^-4^, and 5×10^-5^, respectively).

**Figure S16.**
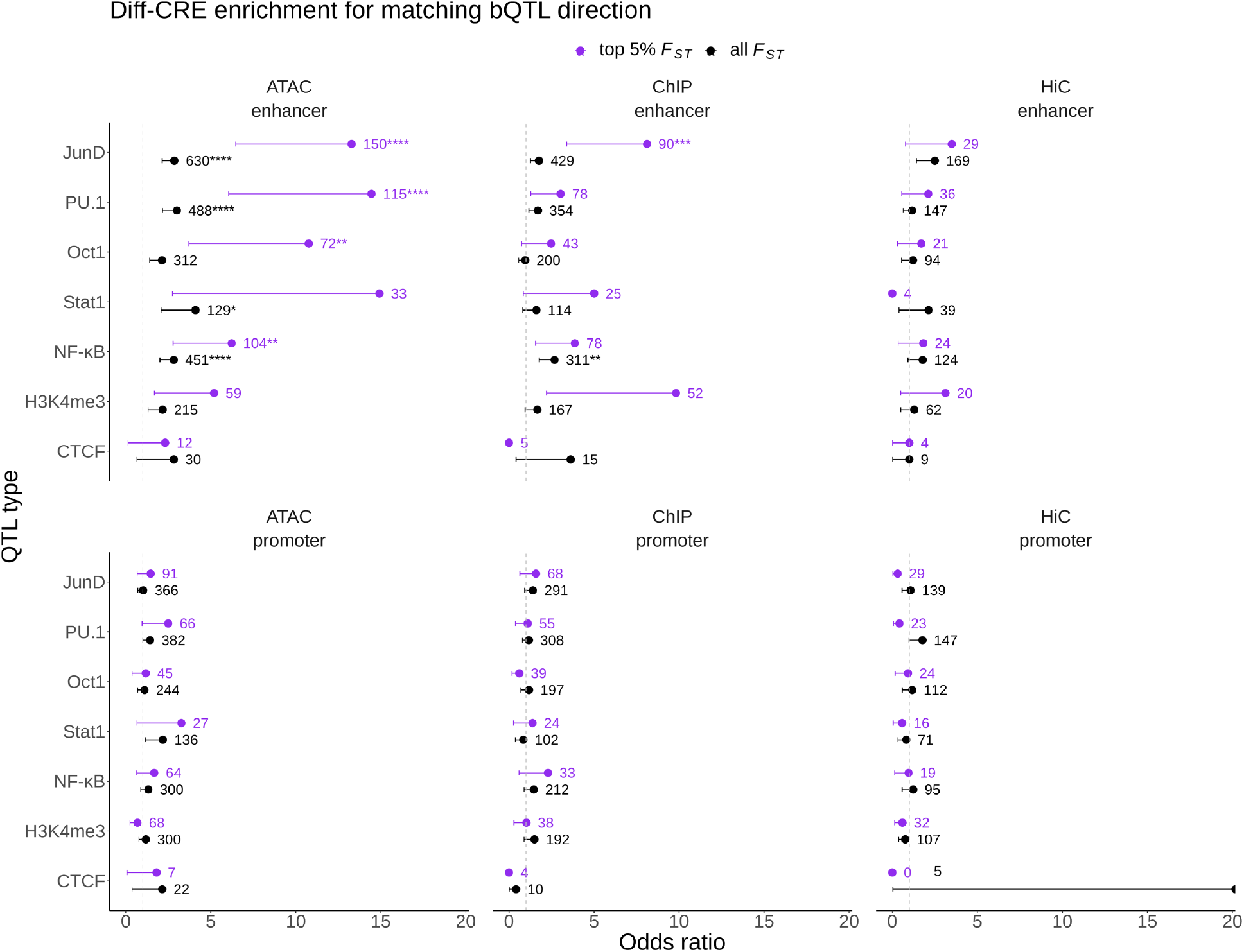
Enrichments for direction matching between bQTL and diff-CREs are driven by enhancers. See Fig. 3e legend.

**Figure S17.**
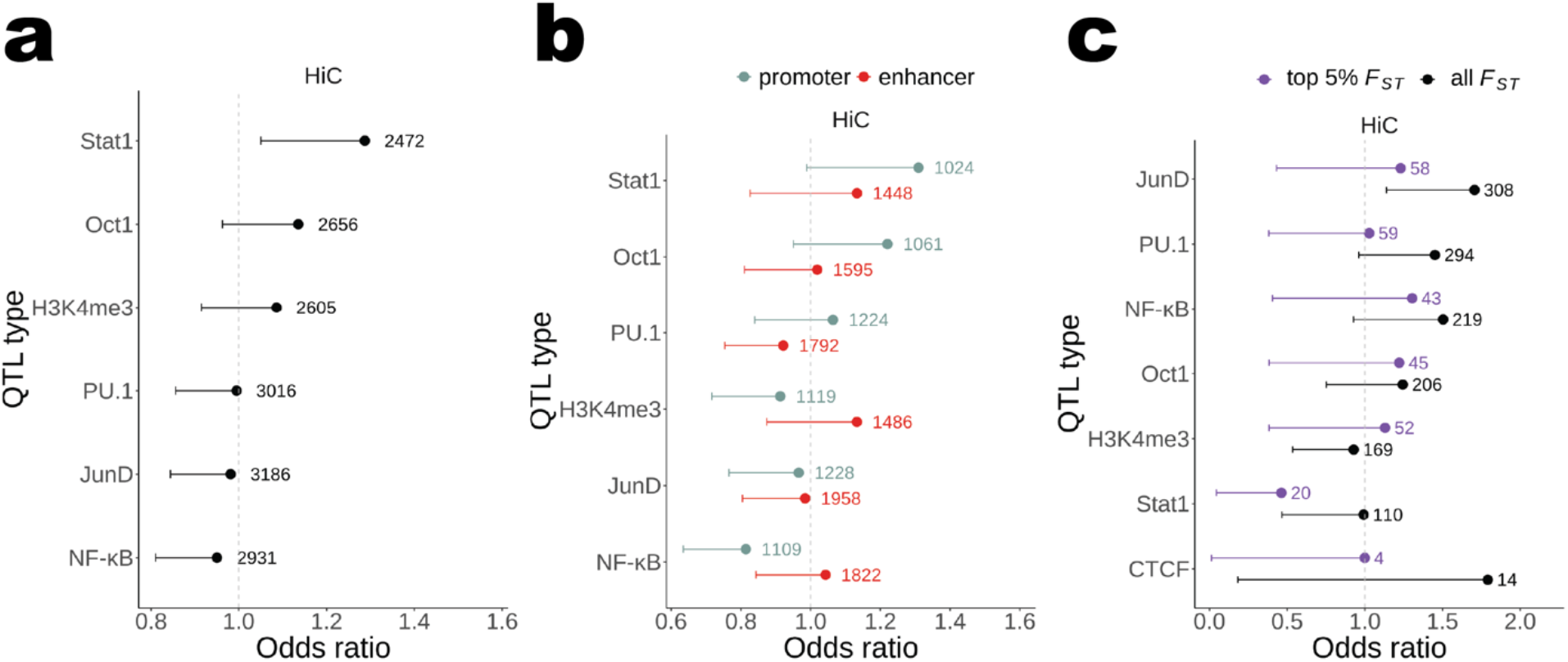
Nominal diff-HiC bQTL enrichment test results. **a)** See Fig. 3b legend. **b)** See Fig. 3c legend. **c)** See Fig. 3e legend.

**Figure S18.**
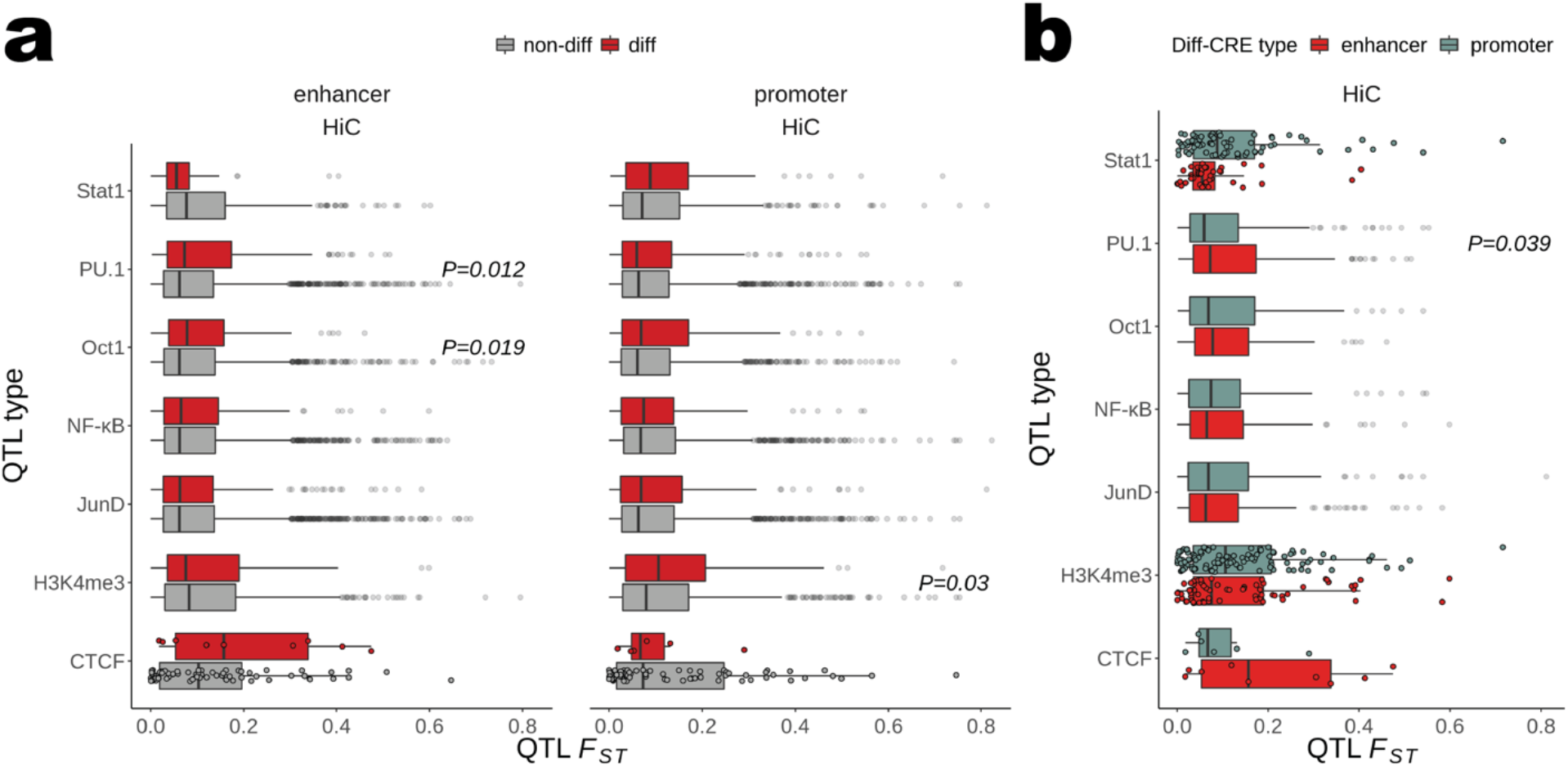
Nominal diff-HiC bQTL ancestry divergence in enhancers versus promoters. **a)** See Fig. 4a legend. **b)** See Fig. 4b legend.

**Figure S19.**
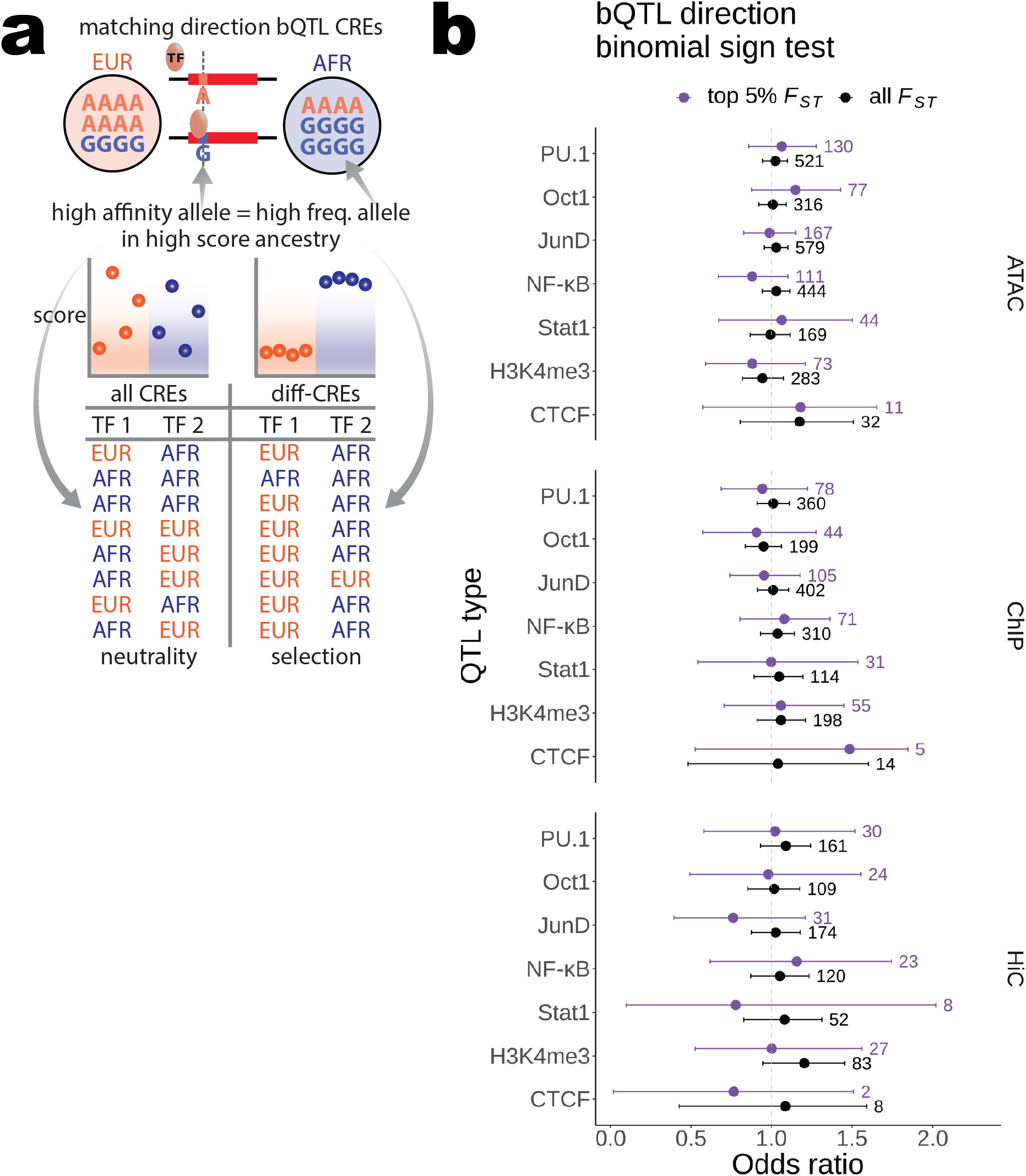
Binomial sign test results on matching direction diff-CRE bQTL. **a)** Schematic illustrating how the binomial sign test (results in (b)) was conducted. The hypothetical G allele at the top is the high affinity bQTL allele and is at higher frequency in AFR, which is also the ancestry with the higher ATAC, ChIP, or HiC scores for the CRE containing this bQTL. This matching direction bQTL CRE contributes one to the “AFR” count for “TF 2” in diff-CREs, whose binding sites therein are under directional selection for greater binding in AFR and/or reduced binding in EUR relative to the proportion of AFR matching direction bQTL CREs genome-wide (expectation under neutrality). **b)** Results of the two-sided binomial test described in (a) are plotted for each QTL type as odds ratios with error bars representing 95% confidence intervals. The total number of diff-CRE bQTL used in each test is shown to the right of each upper bound. None of the P-values pass multiple test correction, so no asterisks are displayed. Binomial test results were normalized to the background probability of success for visualization such that an odds ratio of one represents the background probability of success. Values greater than one indicate the TF has a greater proportion than expected of matching direction diff-CRE bQTL favoring greater binding in AFR.

**Figure S20.**
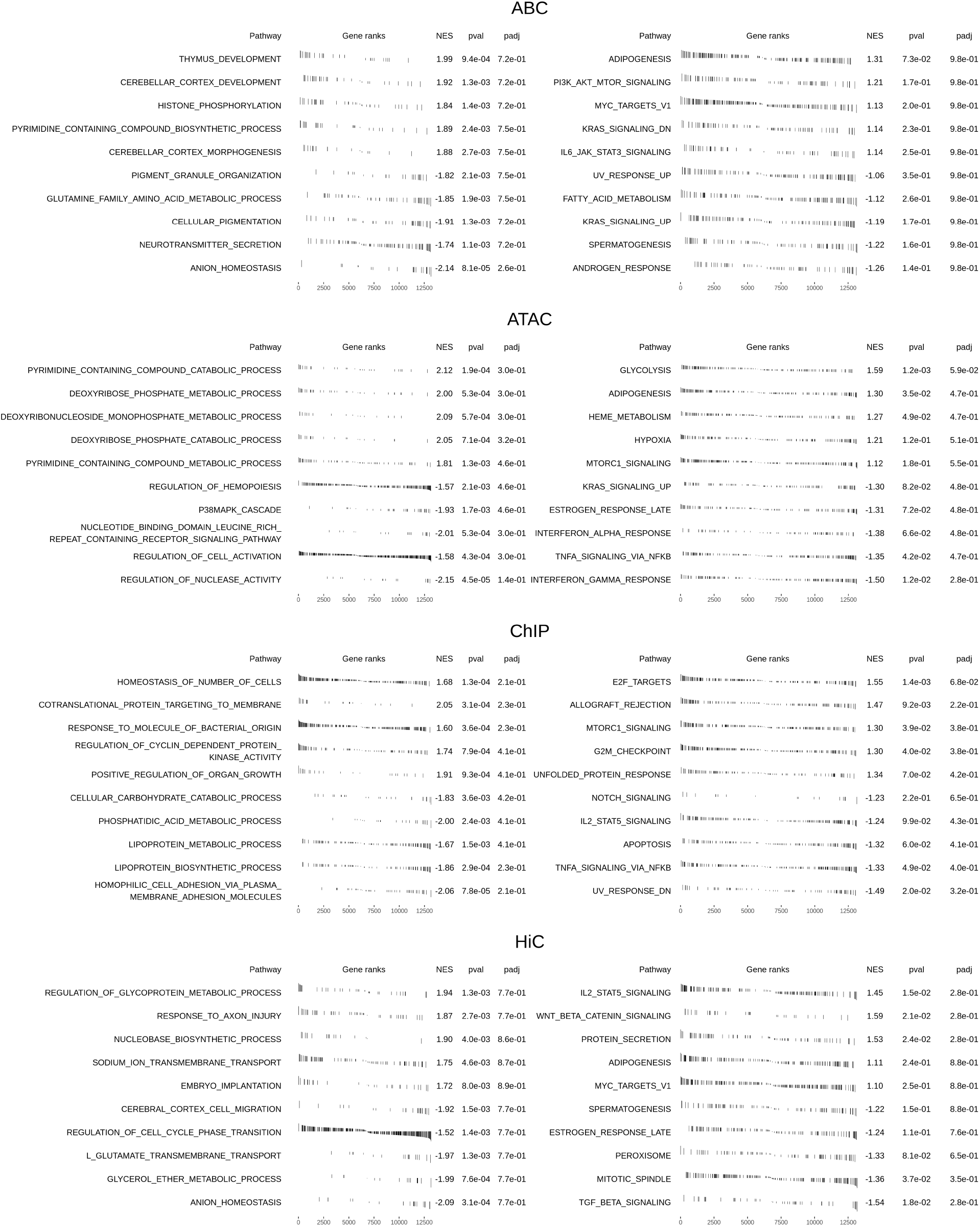
Diff-CRE target gene directional pathway enrichment test results. Results of gene set enrichment analyses from GO biological processes (left) and MSigDB Hallmark gene sets (right) are shown for each score type. A ranking statistic (see Methods), was used to rank all target genes by their most differential CRE from high EUR score to high AFR score. The top 5 processes or gene sets most enriched at the EUR (positive NES) and AFR (negative NES) ends of each list are displayed. Vertical black bars represent the value of the ranking statistic and location in the ranked list where a gene is in a given gene set. Abbreviations: pval = enrichment P-value, NES = normalized enrichment score, padj = Benjamini-Hochberg-adjusted P-value.

### Supplemental tables

Supplemental tables are available at https://doi.org/10.5281/zenodo.7352848. Titles and descriptions of each are below:

- Table S1. tab-separated values (“.txt”). HiChIP read mapping statistics. Number of reads of each category at each mapping step (percentages are of the total read pairs entering each mapping step).
- Table S2. G-zipped, tab-separated values (“.txt.gz”). CEU scores. ABC, ATAC, ChIP, and HiC scores for both CEU replicates for all E-G pairs after filtering across all 16 samples.
- Table S3. G-zipped, tab-separated values (“.txt.gz”). AFR scores. ABC, ATAC, ChIP, and HiC scores for all African ancestry samples for all E-G pairs after filtering across the 14 samples not including CEU.
- Table S4. G-zipped, tab-separated values (“.txt.gz”). EUR scores. ABC, ATAC, ChIP, and HiC scores for all European ancestry samples for all E-G pairs after filtering across the 14 samples not including CEU.
- Table S5. G-zipped, tab-separated values (“.txt.gz”). Differential score statistics. Results of tests for differential ABC, ATAC, ChIP, and HiC scores between AFR and EUR populations for all E-G pairs after filtering across the 14 samples not including CEU.

